# Homeostatic cytokines reciprocally modulate the emergence of prenatal effector PLZF+CD4+ T cells in humans

**DOI:** 10.1101/2022.09.19.508328

**Authors:** Veronica Locher, Sara Park, Daniel Bunis, Stephanie Makredes, Trevor D. Burt, Gabriela K. Fragiadakis, Joanna Halkias

**Author notes:** Corresponding author Address correspondence to: Joanna Halkias, Division of Neonatology, Department of Pediatrics, and Eli and Edythe Broad Center of Regeneration Medicine and Stem Cell Research, UCSF, San Francisco, CA, USA. Current institution (Veronica Locher, Trevor D. Burt).

## Abstract

The development of human adaptive immunity progresses faster than previously appreciated, with the emergence of memory CD4^+^ T cells alongside regulatory T (Treg) cells by the second trimester of pregnancy. We previously identified a prenatal-specific subset of PLZF^+^ CD4^+^ T cells with heightened effector potential that accounted for most memory T cells in the developing intestine and accumulated in the cord blood of infants exposed to prenatal inflammatory pathologies. However, the signals that drive their tissue distribution and effector maturation are unknown. In this report, we define the transcriptional and functional heterogeneity of prenatal PLZF^+^ CD4^+^ T cells and identify compartmentalization of Th-like effector function across the small intestine (SI) and mesenteric lymph nodes (MLN). We find that IL-7, which is more abundant in the SI relative to the MLN, drives the preferential expansion of naïve PLZF^+^ CD4^+^ T cells via JAK/STAT and MEK/ERK signaling. Exposure to IL-7 induces a subset of PLZF^+^ CD4^+^ T cells to acquire a memory-phenotype and rapid effector function, identifying the human analogue of memory-phenotype CD4^+^ T cells. Further, IL-7 modulates the differentiation of Th1- and Th17-like PLZF^+^ CD4^+^ T cells, and thus likely contributes to the anatomic compartmentalization of prenatal CD4^+^ T cell effector function.

## Introduction

Pregnancy is a critical period of early human immune development. Unlike mice, humans are born with a complete set of immune cells, and T cells are evident in the periphery by 12-14 weeks of gestation. Specifically, αβ and γδ T cells emerge simultaneously during prenatal thymopoiesis, and along with regulatory T (Treg) cells, are evident at mucosal surfaces by the second trimester of pregnancy (1–4). Thus, in humans, the critical early life window of development in which the immune system is highly receptive to environmental instruction (5), begins *in utero*.

Growing evidence supports the notion that prenatal T cells are a specialized population adapted to the unique demands of the perinatal period (6). These include the need for tolerance to self and maternal antigens *in utero* and a simultaneous requirement for the generation of a protective immune compartment to face the multitude of environmental antigens encountered after birth. Thus, the survival of a healthy newborn requires the concurrent generation of both regulatory and protective T cells, adding a layer of complexity to the development of human adaptive immunity.

The priming and differentiation of naïve T cells upon encounter with cognate antigen in the context of MHC, co-stimulatory signals, and specific cytokine milieux results in the generation of distinct T helper (Th) cell subsets with characteristic transcriptional and cytokine production profiles (7). In addition to foreign-antigen specific memory Th cells, murine studies have identified memory-phenotype CD4^+^ T cells which arise in response to interactions with self-peptide MHC in combination with homeostatic cytokines in the setting of lymphopenia-induced proliferation (8,9). A recent study identified that early life CD8^+^ T cells have a cell-intrinsic propensity to become virtual-memory cells and constitute the early effector response to infection in adulthood (10). Thus, the human *in utero* environment with limited yet impending exposure to a multitude of foreign antigens after birth, and a peripheral compartment largely void of immune cells, is ideally positioned for the generation of memory-phenotype cells to provide rapid protection at mucosal surfaces. However, whether these cells are present in humans is not clear.

The majority of T cells in mice and humans alike reside in lymphoid and mucosal tissues, with a heterogeneous distribution of subsets and function across specific sites (11,12). In the mature immune system, the anatomic location of T cells is critical to their differentiation and function. It is likely that the developing prenatal immune system similarly integrates local cues to induce the differentiation of cell types tailored to meet the needs of the tissue they reside in. Naïve T cells predominate in most infant and pediatric tissues, yet memory-phenotype CD4^+^ T cells with rapid effector function are evident in the SI by the second trimester of pregnancy (13–16). We recently demonstrated that the majority of these intestinal memory T cells express the transcription factor promyelocytic leukemia zinc finger (PLZF), and though present in lymphoid and non-lymphoid tissues, prenatal PLZF^+^ CD4^+^ T cells specifically accumulated in the intestine and were superior producers of Th1 cytokines as compared to PLZF^-^ CD4^+^ T cells (14). Moreover, PLZF^+^ CD4^+^ T cells were enriched in the cord blood of infants exposed to prenatal inflammatory pathologies, underscoring the clinical relevance of understanding the signals that drive the maturation and compartmentalization of these effector T cells *in utero*.

To better understand how the prenatal environment shapes the developing human CD4^+^ T cell compartment, we set out to identify the cues that drive T cell maturation and function during normal development. We provide evidence of the functional heterogeneity and spatial segregation of memory PLZF^+^ CD4^+^ T cells between the lamina propria of the SI and the draining MLN, revealing a diversity of Th-like effector function that parallels that of conventional CD4^+^ T cells. We further define tissue-specific differences in homeostatic cytokines and identify IL-7 as a critical driver in the distribution and functional maturation of prenatal effector PLZF^+^ CD4^+^ T cells, including the generation of memory-phenotype CD4^+^ T cells in response to lymphopenia-induced proliferation.

## Results

### Prenatal PLZF^+^ CD4^+^ T cells are functionally heterogeneous and display spatially compartmentalized effector function

We previously demonstrated that PLZF^+^ CD4^+^ T cells account for the majority of memory T cells in the developing human SI (14). Moreover, PLZF^+^ CD4^+^ T cells were significantly more likely to produce Th1 cytokines (IFNγ, TNFα) than conventional memory CD4^+^ T cells, displaying a heightened effector potential and a transcriptional signature enriched for T cell activation genes. To determine their range of effector function, we sort-purified intestinal PLZF^+^ CD4^+^ T cells as previously described (14) and analyzed their transcriptional response to PMA/Ionomycin stimulation. In addition to Th1-associated transcripts such as *TNF, IFNG*, and *XCL2*, stimulated PLZF^+^ CD4^+^ T cells also up-regulated transcription of a multitude of cytokines not classically produced by a single T cell subset. These included *IL4* and *IL13*, as well as *IL22* and *LIF* (Figure 1A), suggesting a potentially heterogeneous T cell population composed of distinct effector subsets. To further characterize these effector subsets, we performed single cell RNA sequencing on sort purified PLZF^+^ CD4^+^ T cell from 3 individual donors. Unbiased clustering analysis divided 23,676 cells into 7 subsets represented in each donor and annotated by distinct expression signatures (Figure 1B-E; Supplemental Figure 1A). A subset of Th1-like cells with high expression of *XCL1, XCL2, CCL5, IFNG*, and *CXCR3*, as well as activation (*CD38, CD44, 4-1BB/TNFRSF9* and *NR4A2*) and cytotoxicity (*GZMA, GZMK*, and *NCR3/NKp30*) markers was readily identified. In line with a Th1-like profile, pathway analysis revealed enrichment of IL-12 signaling, T cell receptor (TCR) signaling, and Natural Killer cell mediated cytotoxicity (Figure 1E and F). Transcripts associated with Th17 cells (*CCR6, RORC, RORA, MAF*, and *IL23R*) marked a second subset with increased expression of IL-2, as well as pathway enrichment for both IL-12 and TGFβ signaling. A Th2-like cluster of PLZF^+^ CD4^+^ T cells was characterized by the lineage defining transcription factor *GATA3* and increased expression of *IL32, IL16*, and *MIF*, as well as the TCR signaling regulators *CD5* and *CD6*. Proliferating cells were identified based on higher expression of transcripts related to G1/S phase (*CDC45, PCNA, TYMS*) or G2M phase (*CDK1, TOP2A*) (17), consistent with our previous identification of a large proportion of proliferating memory T cells in the prenatal intestine (14). An interferon-stimulated gene (ISG) signature defined another subset of PLZF^+^ CD4^+^ T cells, which included the higher expression of *DDX58, OAS1, MX1, IFIT1, RSAD2*, and *ISG15*, and was specifically enriched for Type I and Type II Interferon signaling. The ISG signature has been proposed to represent an intermediate Th1 activation state (18) and has also been associated with bystander activation of T cells (19), a defining feature of prenatal PLZF^+^ CD4^+^ T cells. Interestingly, cytokine signaling, T cell activation, and T cell differentiation pathways were enriched among most subsets, whereas TCR signaling was uniquely enriched in Th1-like cells. In stark contrast, Cluster 0 harbored only 22 differentially upregulated genes and was void of cytokine- and TCR-signaling pathways and markers of lymphocyte activation and differentiation (Figure 1D-F), suggestive of a resting or precursor cell state. Cluster 5, while lacking a distinct gene signature, displayed transcriptional overlap with that of the Th-17 like subset, suggestive of an intermediate or transitional state (Figure 1D and E). Lastly, expression of *TNF* and *LIF* transcripts was evident across all PLZF^+^ CD4^+^ T cells without differential expression between subsets (Figure 1G). These distinct transcriptional signatures suggest the presence of different effector branches of prenatal PLZF^+^ T cells that parallel conventional T helper subsets.

**Figure 1.**
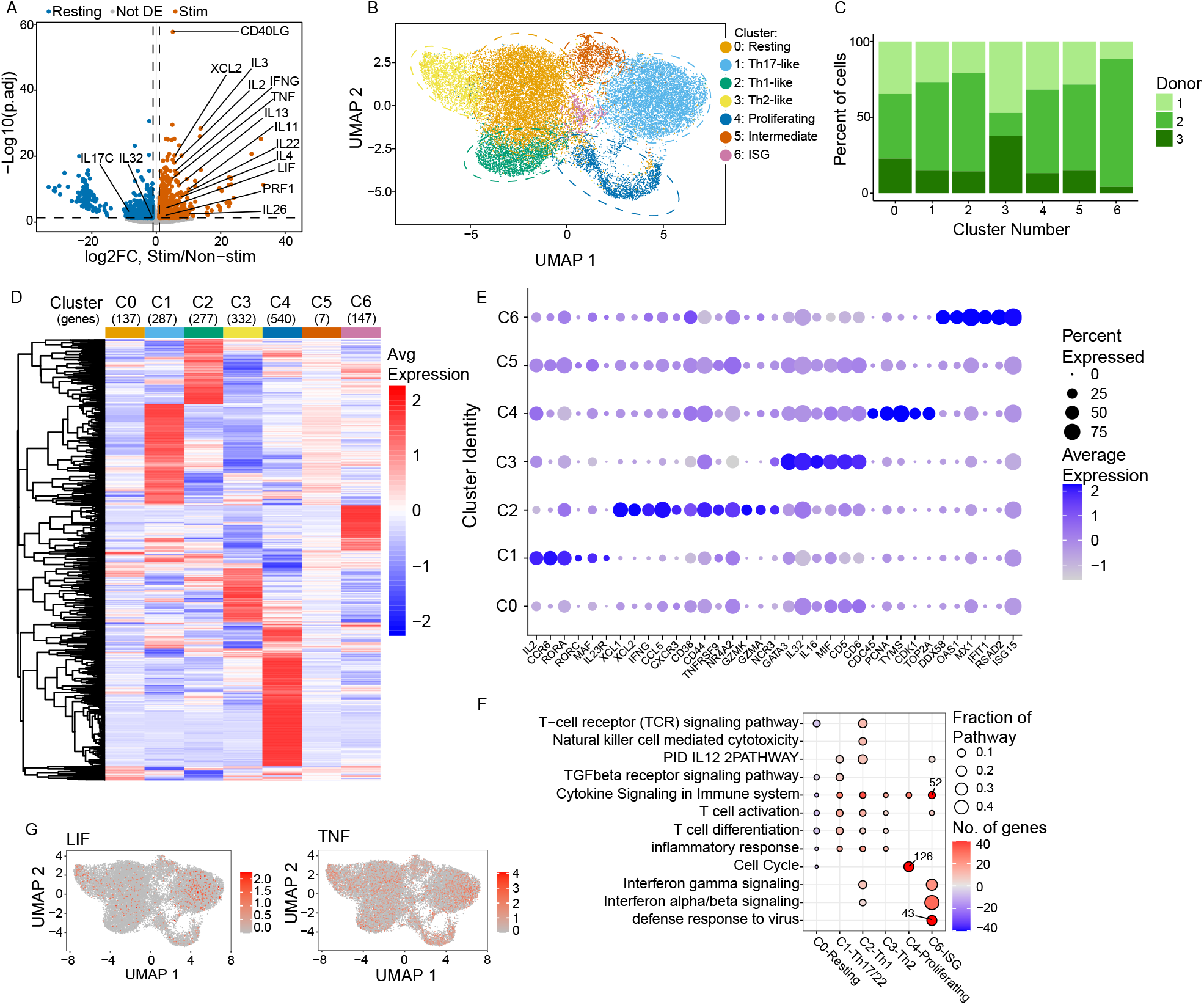
Prenatal PLZF^+^ CD4^+^ T cells in the small intestine are a heterogeneous population. (A) Volcano plot of differentially expressed (DE) genes (log2 fold change > 1, FDR ≤ 0.05) from bulk RNA-seq in stimulated (PMA/ionomycin) versus unstimulated small intestine (SI) PLZF^+^ CD4^+^ T cells identifies enriched (orange) versus diminished (blue) transcripts in stimulated cells. Select cytokines are labeled. (B) UMAP plot of unsupervised (Louvain) clustering of scRNA-seq transcriptional profile of 23,676 sort-purified, PLZF^+^ Vα7.2^-^ CD4^+^ TCRαβ^+^ T cells from the SI of 3 donors, each cluster surrounded by a median centered ellipse and (C) the proportion of cells from each donor in each cluster. (D) Heat map of unsupervised hierarchical clustering of DE genes and (E) dot plot of selected DE genes within each identified cluster of PLZF^+^CD4^+^ T cells. (F) Selected GO terms, Reactome Gene Sets, and KEGG pathways from Metascape analysis of cluster-specific DE gene lists. Downregulated pathways are indicated with negative number of genes and outliers indicated with text overlay. G) UMAP plots of *LIF* and *TNF* expression among SI PLZF^+^CD4^+^ T cells.

We next sought to confirm cytokine production from our predicted Th-like PLZF^+^ CD4 T cell subsets. We previously showed that PLZF^+^ CD4^+^ T cells responded to both TCR activation and to bystander activation in response to cytokines. A role for antigen-independent activation of CD8^+^ T cells has been described in the setting of inflammatory and/or infectious pathologies, yet less is known about bystander activation of human CD4^+^ T cells (20–22). To assess the prenatal Th-like effector program, we stimulated intestinal memory CD4^+^ T cells, with αCD3/CD28 monoclonal antibodies and/or a combination of cytokines known to elicit specific Th-type bystander responses (23). Consistent with our previous findings (14), we observed significantly higher frequencies of IFNγ-producing cells in response to bystander activation among PLZF^+^ CD4^+^ CD45RO^+^ T cells compared to their conventional (i.e., PLZF^-^ CD4^+^) counterparts. Differences in IFNγ-production were also evident by anatomic compartment, with significantly higher proportions of both CD4^+^ memory T cells producing IFNγ in the SI as compared to the MLN, whereas no differences were evident for TNFα (Figure 2A and B, Supplemental Figure 2A). The proportion of IL-13 producing cells in the MLN was higher among PLZF^+^ than PLZF^-^ CD4^+^ CD45RO^+^ T cells, whereas production of IL-4 was similar (Figure 2C-E). Moreover, IL-4 production was mostly absent from the SI, suggesting Th2-like cells in the developing intestine are primarily IL-13 producing cells with minimal capacity for IL-4 production. Despite the presence of ROR γ t^+^ PLZF^+^ CD4^+^ T cells in both the MLN and the SI (Supplemental Figure 2B), IL-17 producing cells revealed the most striking segregation of effector function. While the proportion of IL-17 producing cells was similar among memory CD4^+^ T cells in the MLN across activation conditions, these were essentially absent from the SI (Figure 2F and H). On the other hand, production of IL-22 was significantly higher among memory CD4^+^ T cells of the SI as compared to the MLN and was elicited almost exclusively in response to bystander activation. Further, IL-22 producing cells were significantly more abundant among PLZF^+^ than conventional CD4^+^ memory T cells in the MLN (Figure 2G and H). These findings confirm the presence of functional, cytokine-producing populations that correlate to populations predicted by the transcriptional programs of distinct Th1-, Th2-, Th17-, and Th22-like populations of prenatal PLZF^+^ CD4^+^ T cells and reveal a clear pattern of segregation of effector function between the developing human MLN and SI.

**Figure 2:**
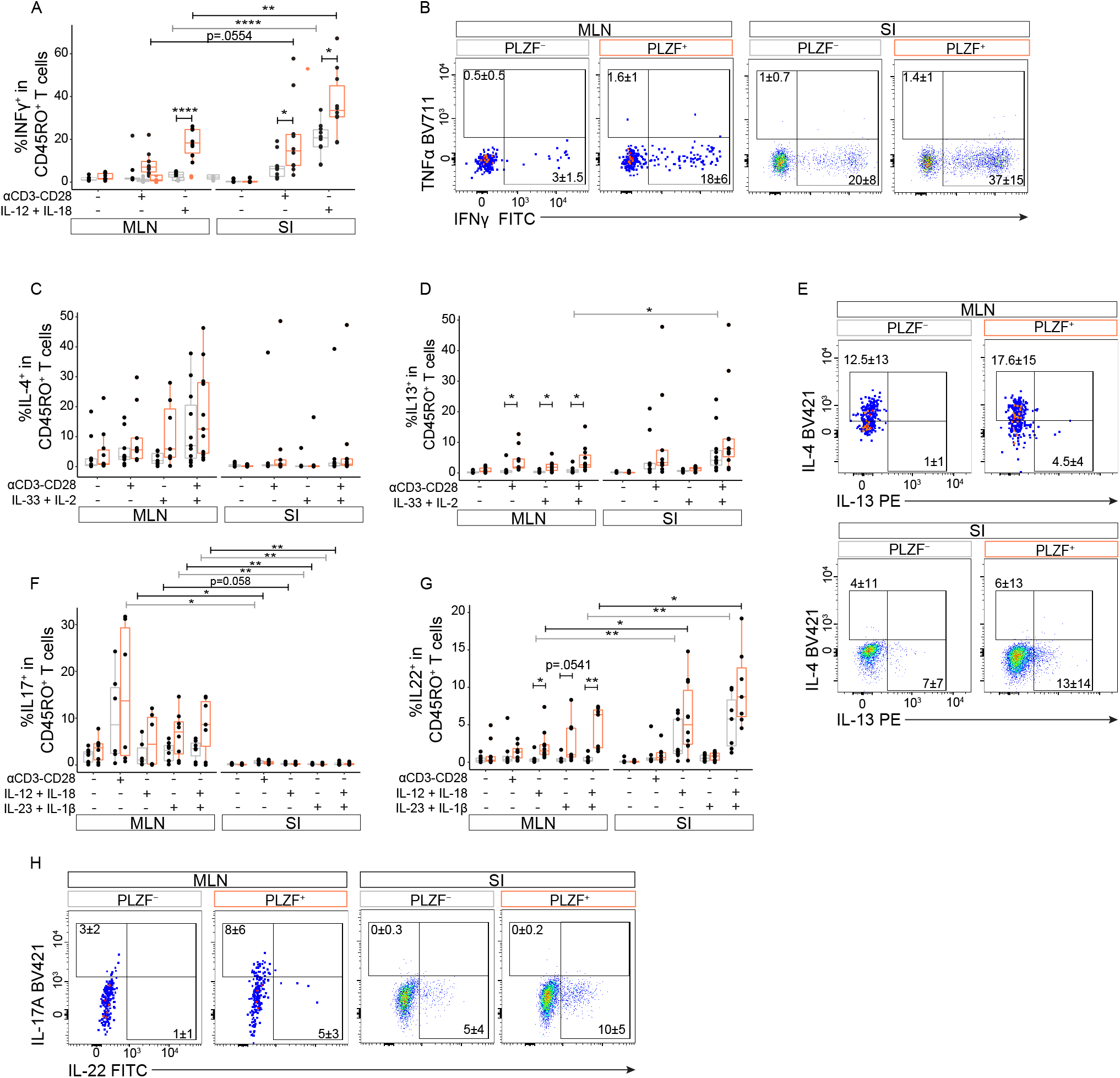
Prenatal PLZF^+^ CD4^+^ T cells display spatially compartmentalized effector function. Quantification (A, C, D, F, G) and representative flow plots (B, E, H) of indicated intracellular cytokine staining after stimulation among PLZF^-^ (grey) and PLZF^+^(orange) CD45RO^+^ CD4^+^ T cells isolated from the small intestine (SI) and mesenteric lymph node (MLN). Cells were stimulated as indicated for 16 hours, and Brefeldin A added in the last 4 hours. Flow plots indicate intracellular cytokine staining following stimulation with IL-12 + IL-18 (B), αCD3/CD28 monoclonal antibodies (mAbs) + IL-33 + IL-2 (D), and IL-23 + IL-1β + IL-12 + IL-18 (H). Circles represent individual donors. Paired ANOVA with Tukey multiple comparison test (A, C, D, F, G) *p < 0.05, **p < 0.01, ***p<0.001 ****p < 0.0001. Large dots used for improved visualization of small cell numbers. Numbers in flow cytometry plots represent means of frequencies of gated populations ± SD.

### IL-7 and TGFβ reciprocally regulate the expansion of prenatal naïve PLZF^+^ CD4^+^ T cells

Based on the intestinal accumulation of PLZF^+^ CD4^+^ T cells (14) and their enrichment for cytokine signaling pathways (Figure 1F), we hypothesized that differences in cytokine levels within tissues might contribute to their disparate distribution. While systemic levels of IL-7 are elevated in the setting of lymphopenia (24), there are significantly higher levels of IL-7 and IL-15 in the prenatal SI as compared to the MLN, while IL-2 was largely undetectable (Figure 3A and Supplemental Figure 3). To examine the influence of these cytokines on naïve CD4^+^ T cells, we opted to study the differentiation of prenatal mature CD4^+^ TCRαβ^+^ CD1α^-^ single positive thymocytes to exclude prior exposure to peripheral homeostatic signals. There was a significant accumulation of PLZF^+^ CD4^+^ T cells in response to IL-7 stimulation in comparison to pre-stimulation (d0) frequencies, an effect that was not evident in the presence of IL-2 or IL-15 (Figure 3B and C). We previously demonstrated that naïve PLZF^+^ CD4^+^ T cells have a 3-fold higher rate of proliferation as compared to their conventional counterparts (14), and here we show that the accumulation in response to IL-7 was driven by the enhanced proliferation of PLZF^+^-relative to that of PLZF^-^ naïve CD4^+^ T cells in a dose-dependent manner (Figure 3D and E).

**Figure 3.**
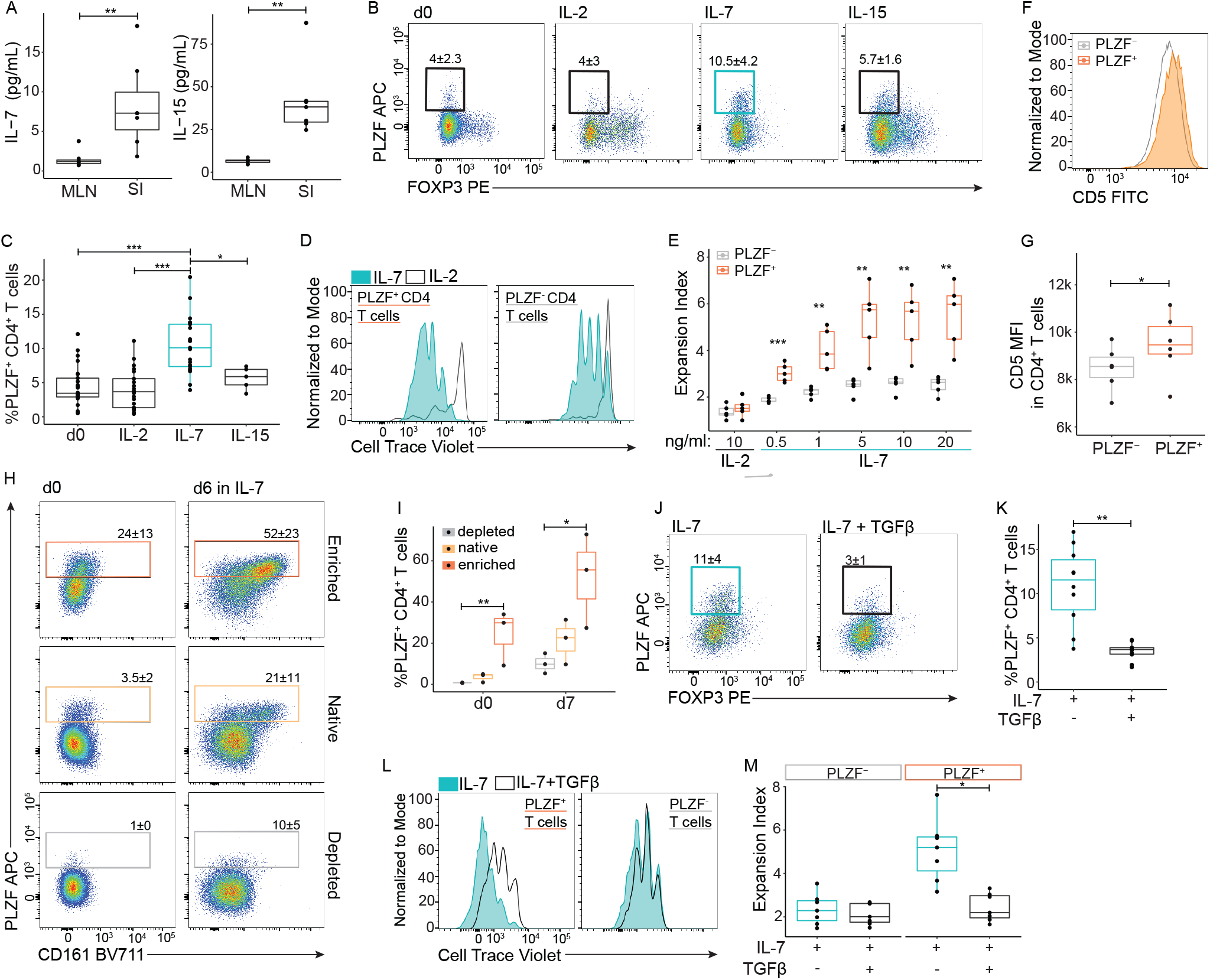
IL-7 and TGFb reciprocally regulate the expansion of prenatal PLZF^+^ CD4^+^ naïve T cells. (A) Quantification of whole tissue concentrations of IL-7 and IL-15 measured by cytokine bead array. (B) Representative flow cytometry plots and (C) frequencies of PLZF^+^ CD4^+^ T cells derived from naïve CD4^+^ T cells before initiation of culture (d0) and after 6 days of culture with IL-2, IL-7, or IL-15 (10ng/mL). (D) Representative histograms of Cell Trace Violet (CTV) dilution, and (E) expansion indices of PLZF^+^ and PLZF^-^ CD4^+^ T cells cultured for 6 days as indicated. (F) Representative histogram and (G) MFI of CD5 expression on PLZF^+^- or PLZF^-^-naïve CD4^+^ T cells. (H) Representative post-sort flow plots and (I) quantification of naïve PLZF^+^ CD4^+^ T cells before initiation of culture (d0) and after 6 days of culture with 10ng/mL IL-7. (J) Representative flow plots and (K) proportion of PLZF^+^ CD4^+^ T cells after 6 days of culture with IL-7 +/- TGFb. (L) Representative histograms of CTV dilution and (M) expansion indices of PLZF^+^ and PLZF^-^ CD4^+^ T cells cultured for 6 days in the presence of IL-7 +/- TGFb (10ng/mL). Circles represent individual donors. Wilcoxon Signed Rank Test (A, G, K, M) and Paired ANOVA with Tukey’s multiple comparison test (C, E, I) *p < 0.05, **p < 0.01, ***p<0.001 ****p < 0.0001. Numbers in flow cytometry plots represent means of frequencies of gated populations ± SD.

Murine studies identify a correlation between surface expression of CD5, a marker of TCR signal strength on naïve CD4^+^ and CD8^+^ T cells, and the intensity of lymphopenia-induced proliferation (25,26). We demonstrate here that naive PLZF^+^ CD4^+^ T cells expressed higher levels of CD5 as compared to their PLZF^-^ counterparts (Figure 3F and G), suggesting that the enhanced response to IL-7 was set during thymic selection (27). To confirm that the effects of IL-7 were due to the expansion of a thymus-derived PLZF^+^ CD4^+^ T population, we sorted on proxy surface markers to enrich and deplete the native population of naïve PLZF^+^ cells (Supplemental Figure 4). This sorting strategy yielded significant differences in the proportion of PLZF^+^ CD4^+^ T cells within the enriched vs depleted fractions which were maintained after 6 days of culture in IL-7 (Figure 3H and I). These data indicate that the accumulation of PLZF^+^ CD4^+^ T cells in response to IL-7 is driven by the enhanced proliferation of a pre-existing population of PLZF^+^ CD4^+^ thymocytes.

While intestinal accumulation of PLZF^+^ CD4^+^ T cells is associated with higher tissue levels of IL-7, reported frequencies of FoxP3^+^ Treg cells are reciprocally increased in the prenatal MLN (13,28) and correlate with elevated levels of TGFβ that contributes to their generation (29). We thus postulated that TGF β would inhibit the accumulation of PLZF^+^ CD4^+^ T cells. Indeed, we found that TGFβ significantly dampened the accumulation of PLZF^+^ CD4^+^ T cells in response to IL-7 by specifically inhibiting their proliferation relative to their PLZF^-^ counterparts (Figure 3J-M). These data provide evidence that variation in steady state cytokine levels likely contributes to the disparate distribution of PLZF^+^ CD4^+^ T cells in prenatal tissues.

### Differences in IL7 signaling contribute to the preferential expansion of PLZF^+^ CD4^+^ T cells

IL-7 is known to drive the homeostatic expansion of human naïve CD4^+^ T cells during early life lymphopenia (30), which led us to determine the basis for the expansion of naïve PLZF^+^ CD4^+^ T cells relative to their PLZF^-^ counterparts. The interaction between IL-7 and the IL-7 receptor (IL-7R/CD127) activates JAK1 and JAK3/STAT5 signaling as well as STAT5-independent signal transduction pathways including PI3K/AKT and MEK/ERK (31). Expression of IL-7R was significantly more prevalent among PLZF^+^ CD4^+^ T cells, whereas expression of the receptors for IL-2 and IL-15 did not differ from those of their PLZF^-^ counterparts (Figure 4A and B and Supplemental Figure 5A-F). Consistent with the pattern of cytokine receptor expression, STAT5B phosphorylation (i.e., pSTAT5B) was induced at higher levels in paired naïve PLZF^+^ as compared to PLZF^-^ CD4^+^ T cells in response to IL-7, but not in response to IL-2 or IL-15 (Figure 4C-D, Supplemental Figure 5G-H). To assess the contribution of STAT5-dependent and - independent signaling in the IL-7 driven expansion of human prenatal PLZF^+^ CD4^+^ T cells, we treated naïve CD4^+^ T cells exposed to IL-7 with a range of signaling inhibitors *in vitro*. As expected, the accumulation of PLZF^+^ CD4^+^ T cells in response to IL-7 was almost entirely dependent on JAK1 and JAK3 signaling. A similar dependency on STAT5 and STAT3 was also evident in response to the combined inhibitor SH-4-54 and was surprisingly replicated by the STAT3-specific small molecule inhibitor STAT3-in-1. We additionally observed significant PLZF^+^ CD4^+^ T cell sensitivity to the selective PI3K inhibitor GDC-0941/Pictilisib and the MEK/ERK pathway inhibitor PD0325901 (Figure 4E). While all signaling inhibitors dampened expansion of PLZF^+^ CD4^+^ T cells, these cells displayed selective inhibition to MEK/ERK signaling which had no discernable effect on their PLZF^-^ counterparts (Figure 4F and G). In addition, there was also a significantly greater effect on the expansion of PLZF^+^ compared to PLZF^-^ CD4^+^ T cells in response to the combined STAT3/5 inhibitor. In sum, the increased frequency of IL7R expression among PLZF^+^ CD4^+^ T cells correlates with higher levels of STAT5B phosphorylation, and there is a selective sensitivity to MEK/ERK signaling in PLZF^+^ CD4^+^ T cells which may additionally contribute to their enhanced proliferation in response to IL-7.

**Figure 4.**
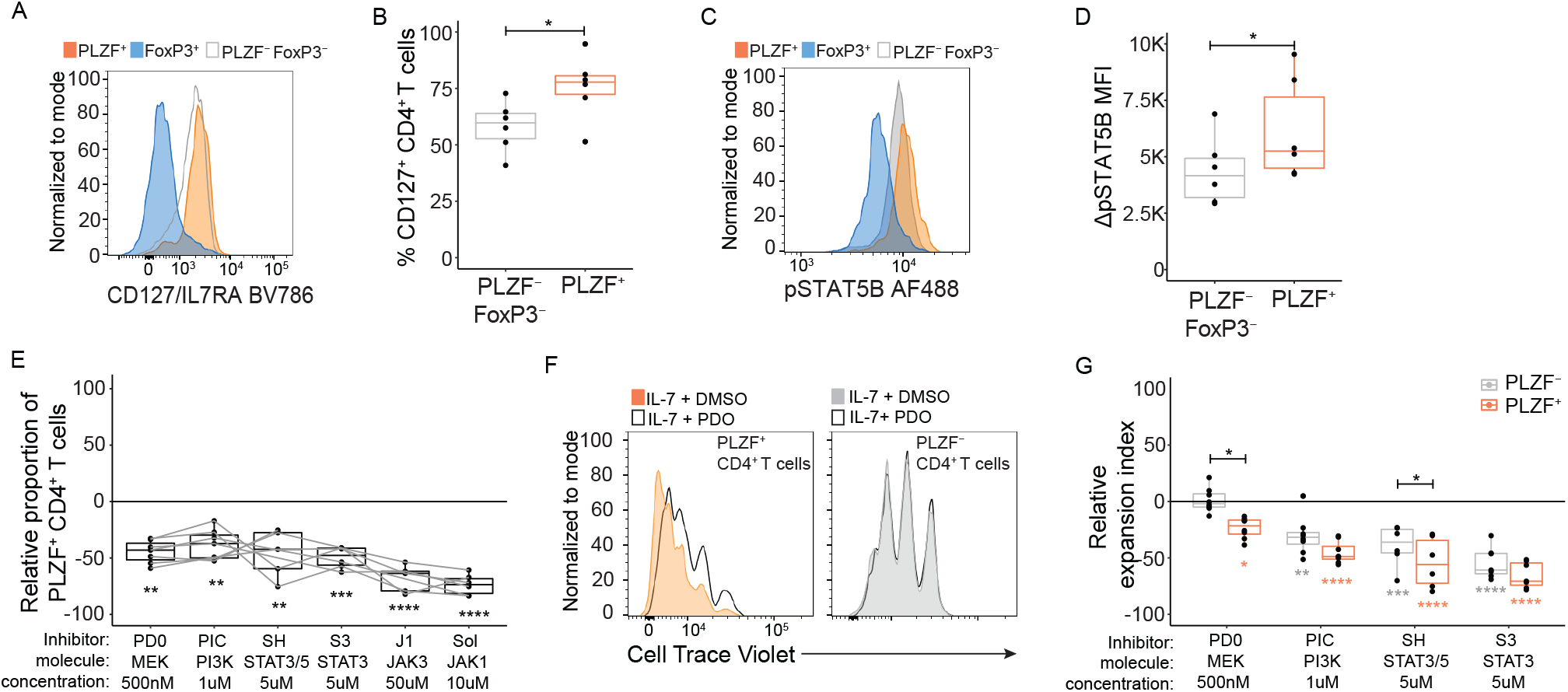
Differences in IL7 signaling contribute to the preferential expansion of PLZF^+^ CD4^+^ T cells. (A) Representative histograms and (B) frequencies of IL7R expression within indicated populations of naïve CD4^+^ T cells. (C) Representative histograms and (D) Normalized MFI of phosphorylated STAT5B (pSTAT5B) among subsets of naïve CD4^+^ T cells after 30 minutes of stimulation with IL-7. (E) Relative proportions of PLZF^+^ CD4^+^ T cells normalized to DMSO control after 6-days of stimulation with IL-7 in the presence of indicated signaling inhibitors. Inhibitors used: PD0 (PD0325901, MEK), PIC (Pictilisib, PI3Kα/δ), SH (SH-4-54, STAT3 and STAT5), S3 (STAT3-IN-1, STAT3), J1 (JANEX-1, JAK3), Sol (Solcitinib, JAK1). (F) Representative histograms of CTV dilution after treatment of naïve CD4^+^ T cells with IL-7 and PD0, and (G) relative increase or decrease in expansion indices among indicated subsets of CD4^+^ T cells after 6 days of stimulation with IL-7 in the presence of indicated signaling inhibitors relative to IL-7 + DMSO control. Circles represent individual donors. Wilcoxon Signed Rank Test (B, D) and Paired ANOVA with Tukey’s multiple comparison test (C, E) *p < 0.05, **p < 0.01, ***p<0.001 ****p < 0.0001. Numbers in flow cytometry plots represent means of frequencies of gated populations ± SD.

### Prenatal PLZF^+^ CD4^+^ T cells are MHC II-restricted and TCR signaling does not interfere with IL-7 driven expansion

IL-7 can function independently of, or synergize with, TCR signaling to promote proliferation of naïve CD4^+^ T cells (32). As prenatal PLZF^+^ CD4^+^ T cells share functional and transcriptional attributes with both conventional and innate-like T cells (14), we next sought to determine whether PLZF^+^ CD4^+^ T cell activation was mediated by recognition of MHC II. We therefore examined the activation of SI memory PLZF^+^ CD4^+^ T cells in the presence or absence of a blocking antibody for MHC class II-TCR interactions during short-term exposure to allogeneic adult CD14^+^ antigen presenting cells *ex vivo*. To quantify the allogeneic response, we calculated the stimulation index, defined as the proportion of the PLZF^+^ CD4^+^ T cells that upregulated expression of the activation molecule CD154 multiplied by the proportion of IFN γ - producing cells (33). We found that the stimulation index of PLZF^+^ CD4^+^ T cells was significantly reduced in the presence of a pan-MHC class II blocking antibody in comparison to isotype control (Figure 5A and B). Thus, consistent with expression of a polyclonal TCR repertoire (14), prenatal PLZF^+^ CD4^+^ T cells recognized antigen in the context of MHC class II, suggesting a stronger similarity to conventional CD4^+^ T cells than to innate-like T cells.

**Figure 5.**
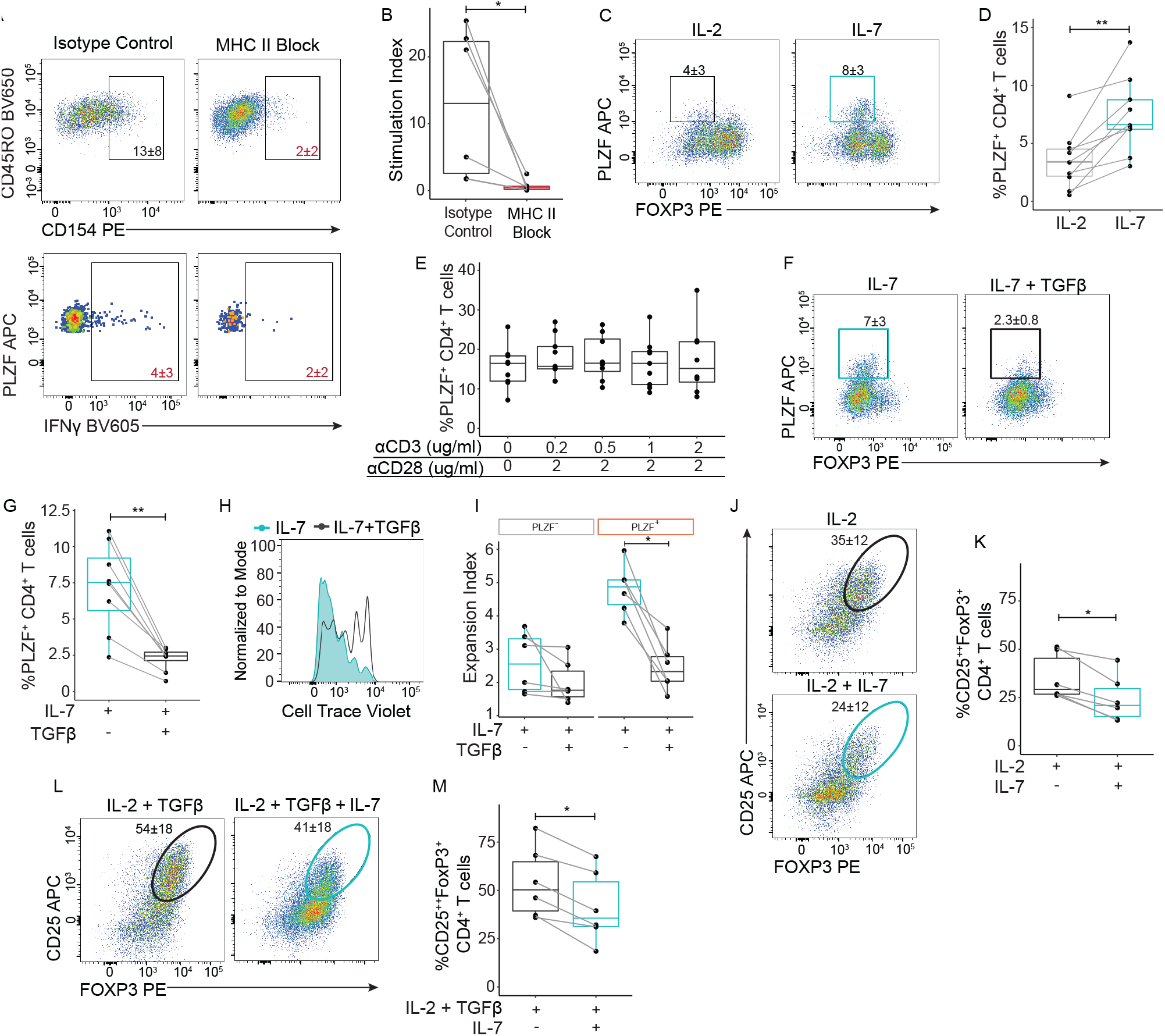
IL-7 and TGFβ sustain reciprocal regulation of MHC Class II restricted PLZF^+^ CD4^+^ naïve T cells in the presence of TCR signaling. (A) Representative flow plots of intracellular CD40L/CD154 expression among SI PLZF^+^CD4^+^ T cells (top) and IFNγ production among CD154^+^PLZF^+^CD4^+^ T cells (bottom) co-cultured with allogeneic adult CD14^+^ antigen presenting cells in the presence of a pan MHC-II blocking antibody (αHLA DR-DP-DQ) or isotype control. (B) Stimulation index, calculated as the percent of CD154^+^ IFNγ^+^ within PLZF^+^ T cells population after 16 hours of co-culture with enriched allogeneic adult CD14^+^ antigen presenting cells in the presence of a pan MHC II blocking antibody (αHLA DR-DP-DQ) or isotype control (IgG2). (C) Representative flow plots and (D) frequencies of PLZF^+^ CD4^+^ T cells derived from naïve CD4^+^ T cells stimulated with αCD3/CD28 in the presence of IL-2 or IL-7 for 6 days. (E) Proportions of PLZF^+^-CD4^+^ T cells stimulated with indicated concentrations of αCD3/CD28 in the presence of IL-7 for 12 days. (F) Representative flow plots and (G) proportions of PLZF^+^ CD4^+^ T cells after 6 days of stimulation with αCD3/CD28 in the presence of IL-7 +/- TGFβ. (H) Representative histograms of CTV dilution, and (I) Expansion indices of PLZF^+^ among indicated CD4^+^T cells after 6 days of stimulation with αCD3/CD28 in the presence of IL-7 +/- TGFβ. (J, L) Representative flow plots and (K, M) frequencies of CD25^hi^ FoxP3^+^ CD4^+^ T cells after 6 days of αCD3/CD28 stimulation in the presence of indicated cytokines. Circles represent individual donors. Wilcoxon Signed Rank Test (B, D, I, K, M) and Paired ANOVA with Tukey’s multiple comparison test (E) *p < 0.05, **p < 0.01, ***p<0.001 ****p < 0.0001. Numbers in flow cytometry plots represent means of frequencies of gated populations ± SD.

We next evaluated the influence of TCR activation on the expansion of PLZF^+^ CD4^+^ T cells in response to IL-7. TCR activation via αCD3/CD28 cross-linking did not affect their preferential accumulation in the presence of IL-7 as compared to IL-2 (Fig. 5C, D), and this was consistent across varying levels of αCD3 stimulation (Fig. 5E). We excluded the possibility that low-level TCR signaling from T cell-T cell interactions (34) might contribute to the IL-7 driven accumulation of PLZF^+^ CD4^+^ T cells as this was unchanged in the presence of a pan-MHC class II blocking antibody (Supplemental Figure 6A and B). Further, TCR engagement had no effect on the ability of TGF β to inhibit the IL-7 driven accumulation and proliferation of PLZF^+^ CD4^+^ T cells (Fig. 5F-I). As prenatal naïve T cells are known to preferentially generate Treg cells we also examined the effect of IL-7 on the differentiation of these (29). Whether naïve CD4^+^ T cells were activated in the absence of Th skewing conditions (Th0; IL-2 alone) or under Treg skewing conditions (IL-2 + TGFβ), addition of IL-7 significantly inhibited the generation of Treg cells (Fig. 5J-M). These data indicate that IL-7 and TGFβ may conversely contribute to the tissue-specific accumulation of PLZF^+^ CD4^+^ T cells and Treg cells.

### Prenatal PLZF^+^ CD4^+^ T cells acquire a memory-phenotype independent of TCR signaling

As long-term polyclonal stimulation of human naïve T cells *in vitro* results in the generation of CD45RO^+^ memory cells (35), we next analyzed induction of a memory phenotype in PLZF^+^ CD4^+^ T cells exposed to IL-7 in the presence of varying levels of CD3/CD28 stimulation. A small proportion of CD45RO^+^ cells emerged at higher levels of prolonged polyclonal stimulation among conventional CD4^+^ T cells, and the frequencies of memory-phenotype cells were significantly higher among PLZF^+^ CD4^+^ T cells across all conditions (Figure 6A and B). Moreover, expression of CD45RO was induced among PLZF^+^ T cells exposed to IL-7 alone and was unaffected by blockade of MHC-TCR interactions (Figure 6B, Supplemental Figure 7A). To determine whether expression of CD45RO corresponded with the potential for cytokine production, we next stimulated CD4^+^ T cells with PMA/Ionomycin after 12 days of culture in the presence of IL-7 and increasing levels of TCR activation. While the potential for TNFα production was high among all subsets of prenatal CD4^+^ T cells, the proportion of TNFα^+^ cells was consistently higher in PLZF^+^ as compared to PLZF^-^ CD4^+^ T cells and did not differ across levels of CD3 activation (Figure 6C and D). Conversely, IFNγ production was nearly absent from PLZF^-^ CD4^+^ T cells, while the proportion of IFNγ^+^ PLZF^+^ T cells was inversely related to the dose of CD3 stimulation (Figure 6C and E). These data indicate that a subset of prenatal PLZF^+^ CD4^+^ T cells acquire a memory-like phenotype and potential for IFNγ production in the absence of TCR stimulation, akin to memory-phenotype murine CD4^+^ T cells.

**Figure 6.**
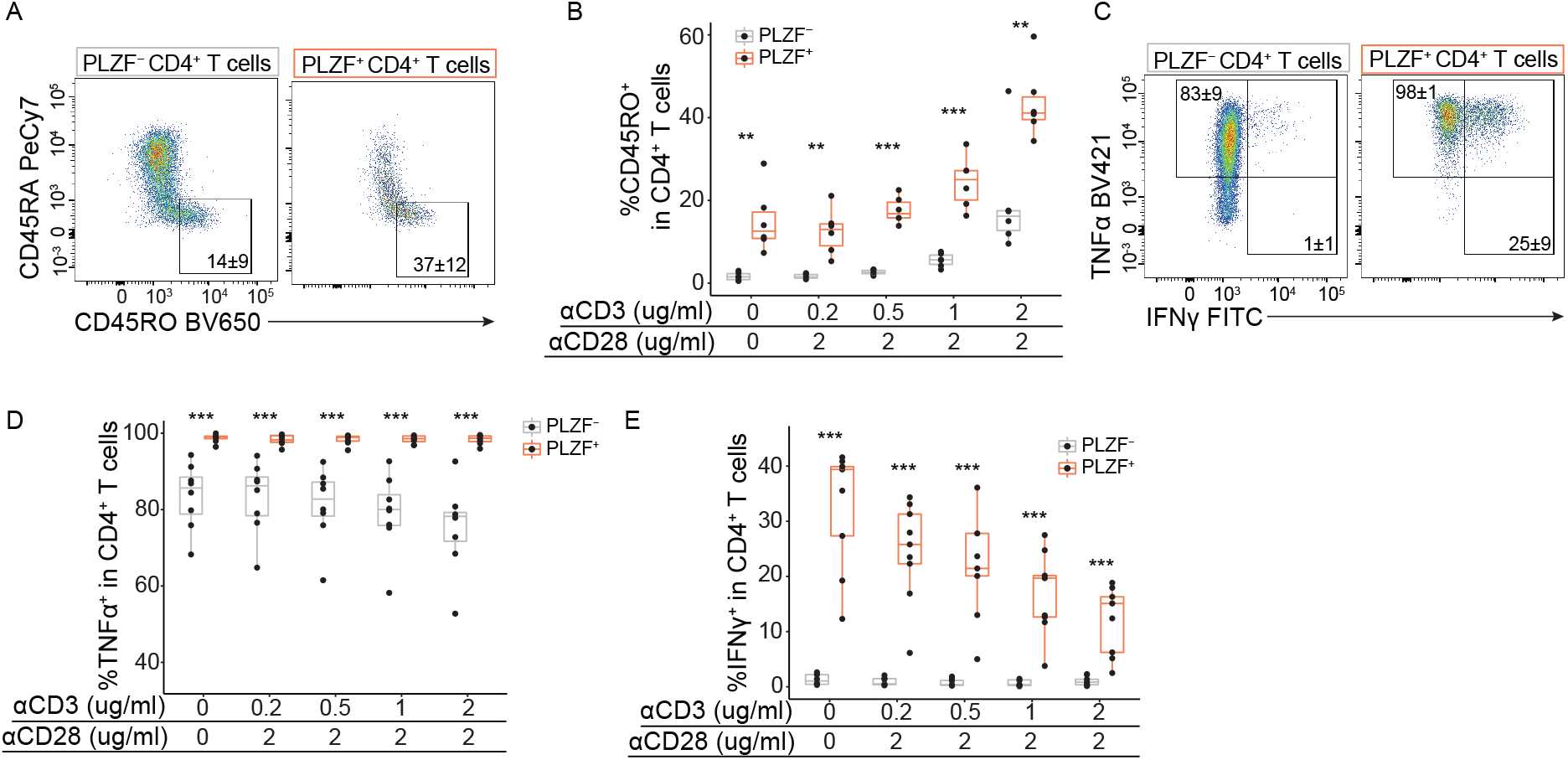
IL-7 alone promotes memory phenotype acquisition in PLZF^+^CD4 ^+^ T cells. (A) Representative flow plots and (B) frequencies of CD45RO^+^ cells among PLZF^+^ and PLZF^-^ CD4^+^T cells after 12 days of culture with IL-7 and indicated αCD3/CD28 concentrations. (C) Representative flow plots of intracellular cytokine staining and (D, E) frequencies of (D) TNF ^+^ cells and (E) IFNγ^+^ cells among CD4^+^T cells after 12 days of culture with IL-7 and indicated αCD3/CD28 concentrations and following re-stimulation with PMA/ionomycin. Circles represent individual donors. Paired ANOVA with Tukey’s multiple comparison test (B, D, E) *p < 0.05, **p < 0.01, ***p<0.001 ****p < 0.0001. Numbers in flow cytometry plots represent mean frequency ± SD of gated populations.

### IL-7 modulates effector function of prenatal PLZF^+^ CD4^+^ T cells

We next asked whether memory-like PLZF^+^ CD4^+^ T cells generated in response to IL-7 alone, or in combination with TCR signaling, could produce cytokines in response to antigen receptor-mediated or bystander activation. To ensure sufficient cell numbers for functional interrogation, we first cultured naïve CD4^+^ T cells in IL-7 for 6 days to promote expansion of PLZF^+^ CD4^+^ T cells and subsequently differentiated them in the presence of αCD3/CD28 and various combinations of cytokines (Supplemental Figure 8A). A subset of PLZF^+^ CD4^+^ T cells produced IFNγ in response to both TCR- and cytokine-mediated activation after prolonged culture in IL-7 alone, as well as TNF in response to TCR activation, while cytokine production was mostly absent from PLZF^-^ CD4^+^ T cells (Figure 7A, Supplemental Figure 8B). Interestingly, differentiation of naïve CD4^+^ T cells in the absence of lineage skewing conditions (Th0) did not generate IFN γ -producing cells in response to TCR-mediated activation yet resulted in significantly higher proportions of PLZF^+^ CD4^+^ T cells capable of IFNγ production in response to bystander activation as compared to their PLZF^-^ counterparts (Figure 7B). Moreover, the proportion of bystander-responsive PLZF^+^ CD4^+^ T cells was further enhanced by addition of IL-7, whereas TNF production was unaffected (Figure 7B, Supplemental Figure 8C). Whole tissue levels of IL-12p70, the bioactive form of IL-12 required for Th1 differentiation, were comparable between the SI and the MLN, indicating that low-level, homeostatic production could contribute to the generation of these effector T cells (Supplemental Figure 8D). Though Th1 differentiation induced the generation of TCR-responsive, IFNγ - and TNF -producing CD4^+^ T cells, the proportion of these was consistently higher among PLZF^+^ CD4^+^ T cells. Further, addition of IL-7 to Th1 differentiation conditions specifically enhanced the proportion of IFNγ - producing PLZF^+^ CD4^+^ T cells in response to both TCR-mediated and bystander activation yet had no effect on their PLZF^-^ counterparts (Figure 7C and D, Supplemental Figure 8E).

**Figure 7.**
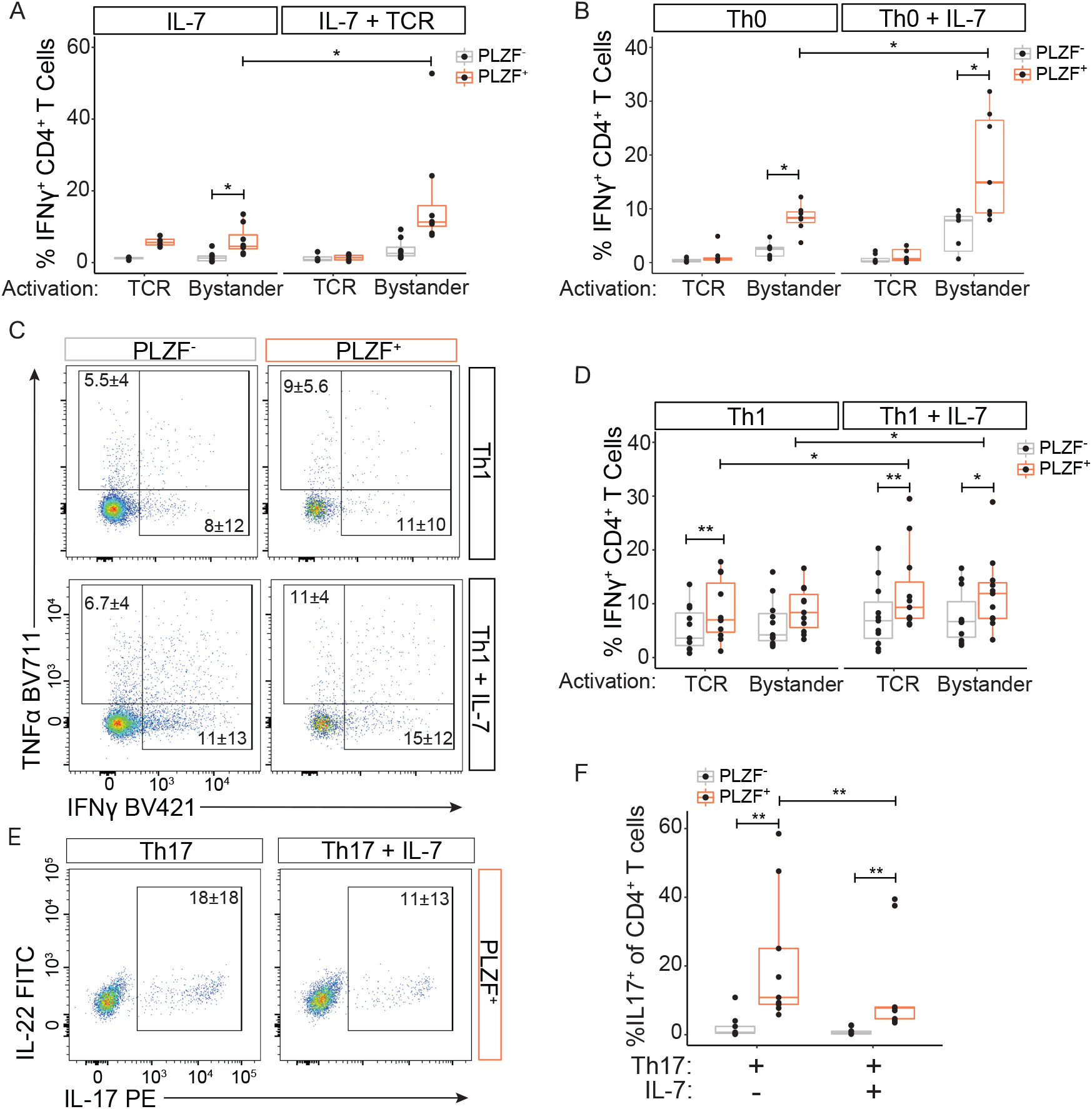
IL-7 modulates the effector maturation of prenatal PLZF^+^ CD4^+^ T cells. Functional studies following a two-step in vitro culture in which naïve CD4^+^ T cells were first expanded in the presence of IL-7 alone, followed by 7 days of maturation in the indicated conditions: IL-7 alone, IL-7 + TCR (stimulated with αCD3/CD28 mAbs), Th0 (TCR + IL-2); Th1 (TCR + IL-12), Th17 (TCR + IL-23 + IL-1β) (A, B, D) Proportions of IFNγ^+^ cells among CD4^+^ T cell subsets matured in the indicated conditions and following 24 hour re-stimulation with αCD3/CD28 (TCR) or IL-12/IL-18 (Bystander). (C) Representative flow plots of intracellular cytokine staining among CD4^+^ T cells differentiated under indicated conditions following re-stimulation with αCD3/CD28. E) Representative flow plots of intracellular cytokine staining and F) proportions of IL-17^+^ cells among indicated CD4^+^ T cells matured under Th17-skewing conditions +/- IL7 and following PMA/ionomycin stimulation. Circles represent individual donors. Wilcoxon Signed Rank Test to compare PLZF^+^ to PLZF^-^ fractions and paired stimulations across maturation conditions (A, B, D, F). Numbers in flow cytometry plots represent means of frequencies of gated populations ± SD.

Despite the observed compartmentalization of IL-17 producing CD4^+^ T cells to the MLN (Figure 2F), tissue levels IL-1β and IL-23 required for human Th17 differentiation were significantly higher in the SI (Supplemental Figure 8F). Naive CD4^+^ T cells differentiated in the presence of IL-1β and IL-23 induced the capacity for IL-17 production almost entirely within the PLZF^+^ subset, and the proportion of these was unaffected by addition of IL-6, or IL-6 plus TGFβ (Supplemental Figure 8G). However, addition of IL-7 to Th17 differentiation conditions significantly dampened the potential for IL-17 production among PLZF^+^ CD4^+^ T cells (Figure 7E, F). Together, these data reveal a role for IL-7 in modulating effector maturation of prenatal PLZF^+^ CD4^+^ T cells, where IL-7 reciprocally enhances production of IFNγ and inhibits the potential for IL-17 production. This divergent maturation in response to IL-7 is likely a contributing factor to the observed compartmentalization of prenatal CD4^+^ T cell effector function between the prenatal SI and the MLN.

## Discussion

This study provides evidence of the emergence of human protective adaptive immunity with anatomically compartmentalized effector function during the second trimester of human gestation that parallels conventional Th1, Th2, Th17, and Th22 cells. Through transcriptional analysis, protein validation, and *in-vitro* culture of primary human cells we identified a critical role for IL-7 in the tissue-specific accumulation and effector maturation of human prenatal PLZF^+^ CD4^+^ T cells that likely contributes to their intestinal segregation. These results build on the current shift in our understanding of human prenatal immunity from one of functional immaturity to that of a specialized adaptation to meet the demand for prenatal tolerance and concurrently provide rapid postnatal protection.

Layering, or progressive transitions in the developmental origin, of CD4^+^ T cells is thought to underlie the preferential generation of Treg cells upon antigen-mediated activation of prenatal naïve T cells in the periphery (29). Our previous work extended our understanding of prenatal adaptive immunity to include human PLZF^+^ CD4^+^ T cells with rapid Th1-like effector function that were absent from the adult (14). Evidence of specialized early life adaptive immunity is well established in mice, in which neonatal CD8^+^ T cells preferentially develop into virtual-memory T cells with innate immune functions and play a role in the early response to infection (10). Less is known about the origin and function of memory-phenotype CD4^+^ T cells, which are thought to exist in humans but have been mostly studied in mice.

In the setting of physiologic early life lymphopenia, the peripheral T cell pool is driven by thymic output and rapid expansion of naïve T cells in an environment with high levels of homeostatic cytokines (36–38). A central role for IL-7 has been identified in the homeostatic proliferation and survival of murine and human naïve CD4^+^ T cells (30,39). Moreover, lymphopenia induced proliferation in neonatal mice resulted in the generation of memory phenotype CD4^+^ T cells with rapid effector function (9). Subsequent studies demonstrated that the rapid proliferation of memory-phenotype CD4^+^ T cells in young mice was driven by IL-7 (40). Systemic levels of IL-7 are increased in lymphopenic hosts (24), with the highest levels reported in lymphoid organs such as the thymus and lymph nodes (41). It is thus notable that we detected significantly higher levels of IL-7 protein in the human prenatal intestine as compared to the draining lymph nodes, which mirrors the preferential accumulation of memory PLZF^+^ CD4^+^ T cells with enhanced Th1-like effector function in the intestine. Our data provide evidence for the IL-7 driven proliferation, and the acquisition of memory-phenotype and innate function among a subset of thymic-imprinted, naïve PLZF^+^ CD4^+^ T cells, and identify the human analogue of early life memory-phenotype CD4^+^ T cells.

In mice, the generation of memory-phenotype CD4^+^ T cells is dependent on tonic, low-level signaling from TCR-MHC class II interactions and thought to be driven primarily in response to self-antigens (8). Self-reactivity is evident among human prenatal CD4^+^ T cells (29), and we and others have also demonstrated that prenatal intestinal CD4^+^ T cells responded to microbial antigens (42,43). Although blocking TCR-MHC class II interactions in the short term did not influence the proliferation or the acquisition of memory-phenotype of PLZF^+^ CD4^+^ T cells *in vitro*, it is possible that TCR signals are required for their long-term survival *in vivo* (44). Future work will determine whether IL-7-expanded and *ex-vivo* isolated prenatal T cells are responsive to self- and/or foreign antigens.

T cell activation in response to cognate antigen remains one of the hallmark features of adaptive immunity. In the setting of early life, recognition of danger signals through antigen-independent activation may serve a rapid protective function at barrier surfaces. Similar to virtual memory CD8^+^ T cells, memory-phenotype CD4^+^ T cells have been reported to provide early protection against pathogens (8,45), suggesting a vital role in the setting of early life immunity. In stark contrast to the mucosal predominance of Th17 cells in mice (46) and human adults (47), prenatal IL-17 producing CD4^+^ T cells segregated to the MLN and were absent from the SI, which instead contained higher frequencies of IL-22 producing cells. The anatomic segregation of steady state prenatal IL-17- and IL-22-producing CD4^+^ T cells to the MLN and SI respectively may serve to limit *in utero* inflammation upon encounter with swallowed luminal antigens or inflammatory mediators while promoting the development and maintenance of epithelial barrier function (48). Production of IL-22 among CD4^+^ T cells was elicited almost exclusively to bystander activation, which suggests a memory-phenotype origin, and more work is needed to identify the cues that drive their effector maturation.

The heterogeneity and spatial segregation of effector function posits that PLZF^+^ CD4^+^ T cells are composed of a combination of lymphopenia-induced, memory-phenotype cells as well as antigen-experienced memory T cells. Though PLZF expression is imprinted during thymic development, neither mature CD4 thymocytes nor naïve peripheral CD4 T cells were capable of IFNγ production, indicating that unlike MAIT or NKT cells, additional steps were required for effector maturation (14). We show that PLZF^+^ CD4^+^ T cells recognize antigen in the context of MHC class II, which coupled with a polyclonal TCR repertoire (14), indicates significant overlap with conventional CD4^+^ T cells, including the presence of distinct Th-like subsets. Migratory antigen presenting cells capable of sensing pathogen and stimulating T cells are evident by the second trimester of gestation (49), and antigen-specific immunity to prenatal pathogen exposure is well documented (50), indicating that Th-differentiation is likely occurring *in utero*. We detected low yet consistent levels of Th-skewing cytokines within the prenatal intestine, which consistently elicited a more robust effector response from PLZF^+^ CD4^+^ T cells *in vitro*. Moreover, our data identify a role for IL-7 in the modulation of Th1- and Th17-like differentiation among PLZF^+^ CD4^+^ T cells and provide a plausible mechanism for the tissue-specific segregation of prenatal effector function.

Beyond extending our understanding of human immune ontogeny, an improved understanding of the composition, spatial distribution, and functional responses of prenatal T cells is foundational to the development of early life immunomodulatory interventions with potentially lifelong consequences on immune health.

## Methods

### Cell Isolation

PBMCs from adult blood were isolated by Ficoll-Histopaque (Sigma-Aldrich) gradient centrifugation and cryopreserved in freezing medium (90% FBS + 10% DMSO; ATCC) for batch analysis. Prenatal organs were processed within 2 hours of collection as previously described (14). Briefly, the MLN and SI were dissected, washed, and cut into 1 cm fragments. Mucus from the SI was removed by DTT washes, and the SI epithelial layer was removed by EDTA washes. The MLN and SI were digested in collagenase IV (Life Technologies) and DNase (Roche), filtered, and lymphocytes isolated by 20/40/80 Percoll (GE Healthcare) gradient centrifugation. Intestinal CD4 T cells were isolated by negative selection using the EasySep Human CD4 T Cell Isolation Kit (STEMCELL Technologies). Thymocyte single cell suspensions were obtained by pressing minced tissue though a 70μm strainer. Mature CD4 single positive (SP) thymocytes were negatively enriched using the EasySep Biotin Positive Selection Kit for depletion (STEMCELL Technologies) as previously described (51). Viability was measured with Trypan Blue (Sigma Aldrich).

### Antibodies and Flow Cytometry

Cells were incubated in 2% FCS in PBS with 1 mM EDTA with human Fc block (STEMCELL Technologies) and stained with Aqua LIVE/DEAD Fixable Dead Cell Stain Kit (Invitrogen) and fluorochrome-conjugated antibodies against surface markers. Intracellular protein and cytokine staining was performed using the Foxp3/Transcription Factor Staining Buffer Set (Tonbo Biosciences). Mouse and rat anti-human monoclonal antibodies (mAbs) used in this study included: IFNγ FITC (clone 25723.11, BD Biosciences, cat# 340449), TCRαβ Percp 710 (clone IP26, Invitrogen, cat# 46-9986-42), TCRγδ Pe-CF594 (clone B1, BD Biosciences, cat# 562511), CD45RA PE-Cy7 (clone HI100, BD Biosciences, cat# 560675), CD4 APC-H7 (clone L200, BD Biosciences, cat# 560837), PLZF APC (clone 6318100, R&D, cat# IC2944A), TCR Va7.2 BV605 (clone 3C10, BioLegend, cat# 351720), CD45RO BV650 (clone UCHL1, BioLegend, cat# 304232), TNFα BV711 (clone MAb11, BioLegend, cat# 502940), CD161 BV785 (clone HP-3G10, BioLegend, cat# 339930), CD45 BUV395 (clone HI30, BD Biosciences, cat# 563792), αCD3 BUV737 (clone UCHT1, BD Biosciences, cat# 612750), CD8 APC-R700 (clone RPA-T8, BD Biosciences, cat# 565166), IL22 FITC (clone 22URT1, Invitrogen, cat# 11-7229-42), IFNg BV421 (clone 4S.B3, BD Biosciences, cat# 564791), IL2 BV711 (clone 5344.111, BD Biosciences, cat# 563946), IL17A BV421 (clone BL168, BioLegend, cat# 512322), IFNγ BV711 (clone 4S.B3, BioLegend, cat# 502539), IL13 PE (clone JES10-5A2, BioLegend, cat# 501903), IL4 BV421 (clone MP4-25D2, BioLegend, cat# 500826), Vα7.2 Biotin (clone 3C10, BioLegend, cat# 351724), Vα2.4 Biotin (clone 6B11, Invitrogen, cat# 13-5806-82), Streptavidin APC-R700 (BD Biosciences, cat# 565144), CD154 PE (clone TRAP1, BD Biosciences, cat# 555700), TNFα PECy7 (clone MAb11, BD Biosciences, cat# 557647), CTV BV421 (Thermo Fisher, cat# C34571), IFNγ BV605 (clone 4S.B3, BioLegend, cat# 502536), CD8 BV711 (clone RPA-T8, BD Biosciences, cat# 563677), CD215 BV421 (clone JM7A4, BD Biosciences, cat# 747704), CD127 BV786 (clone HIL-7R-M21, BD Biosciences, cat# 563324), CD25 FITC (clone M-A251, BioLegend, cat# 356106), CD8 FITC (clone RPA-T8, BioLegend, cat# 301050), CD161 BV711 (clone DX12, BD Biosciences, cat# 563865), FOXP3 PE (clone PCH101, Invitrogen, cat# 12-4776-41), PD-1 BV605 (clone EH12.2H7, BioLegend, cat# 329924), CD8 PeCY7 (clone RPA-T8, BD Biosciences, cat# 557746), Streptavidin BV605 (BioLegend, cat# 405229), CCR7 BV421 (clone G043H7, BioLegend, cat# 353208), CD4 APC-Cy7 (clone RPA-T4, BioLegend, cat# 300517), Aqua LD (Invitrogen, cat# L34957).

For phosphoflow staining, 1M CD4^+^ SP thymocytes were stimulated with IL-7 at 5ng/mL for 30 minutes at room temperature. The reaction was stopped by adding Fix/Perm and subsequently stained for pSTAT5B (pY694, BD Biosciences) in a 96-well plate using the Foxp3/Transcription Factor Staining Buffer Set. All data were acquired with an LSR/Fortessa Dual SORP flow cytometer (BD Biosciences) and analyzed with FlowJo V10.0.8 (TreeStar) software. Expansion Index was calculated using FlowJo software according to the formula: expansion index= (1-PF)/(1-Dil), where PF=fraction of the original population dividing at leads once during the culture period and Dil=percentage of cells in the final population that have divided.

### Bulk RNA seq

T cells were isolated as described above from the intestine of 3 individual samples and PLZF^+^ CD4^+^ TCR-β^+^ T cells were sorted on proxy surface markers using FACSAria2 SORP (BD Biosciences) as previously described (14). Half the sorted cells were stimulated with 50 ng/ml PMA (Santa Cruz Biotechnology Inc.) and 5μg/ml Ionomycin (Sigma-Aldrich) for 3 hours at 37°C in 4% O2, the remaining cells were left unstimulated. RNA was extracted and purified with the Dynabeads mRNA DIRECT Purification Kit (Thermo Fisher Scientific). mRNA libraries were constructed using the Nugen/Nextera XT Library Prep Kit (Illumina), and 3 samples (3 donors) were sequenced on an Illumina HiSeq 4000 by the Functional Genomics Core Facility at UCSF. The reads from the Illumina HiSeq sequencer in fastq format were verified for quality control using the fastqc software package. Reads were aligned to the human genome (hg38) and read counts aggregated by gene using the Ensembl GRCh38.78 annotation using STAR (52). The transcriptome of unstimulated prenatal PLZF^+^ CD4^+^ T cells used for comparison was previously published (14). Differential gene expression analysis was performed on all genes with at least 10 reads with the DESeq2, version 1.16.1 (53). Volcano plots were created with ggplot2 depicting DE genes (log2 fold change > 0.5, FDR ≤ 0.05).

### Single Cell RNA-seq

Single, live PLZF^+^CD4^+^ TCRαβ^+^ T cells from the SI were sorted on proxy markers as previously described (14). Post-sort purity was > 87% as determined by intracellular staining for PLZF. Single cells were captured by droplet-based microfluidics, lysed and sequencing libraries prepared in a single bulk reaction following the 10X Genomics protocol, and transcripts were sequenced using a HiSeq4000 System (Illumina). FASTQ files from each multiplexed capture library were mapped to a custom reference containing GRCh19 using the cellranger (v2.0.0) (10X Genomics) count function. After demultiplexing cells into samples, Seurat (v3.1.5) was used to perform quality control filtering of cells. Cells were retained if they a) contained ≥500 reads, b) ≤5% reads coming from mitochondrial genes, and c) called a singlet by Demuxlet, leaving 23,676 cells for downstream analysis. Data was log normalized then principal components analysis was run on variable features selected by the FindVariableFeatures function after scaling with regression of percent mitochondrial and nReads per cell. UMAP and Louvain clustering were performed with Seurat defaults and resolution 0.3. Differential gene expression between subsets of PLZF^+^ CD4^+^ T cells was performed using FindAllMarkers with Seurat’s MAST implementation (54). Genes were deemed significantly different if an FDR < 0.05, an average fold change > 1.2, and the gene was detected in >5% of cells in either comparison group. Visualizations were created with dittoSeq (v1.6.0) (55). Gene set analyses were performed using Metascape (56) with visualizations created in R with ggplot2.

### CD4 T cell Stimulation assays

Single cell suspensions of isolated CD4 T cells from the SI and MLN were cultured in a 96-well U-bottom plate in complete RPMI (cRPMI) supplemented with 10% FBS, 10 mM HEPES, 2 mM L-glutamine, and 1X nonessential amino acids and stimulated with plate-bound αCD3 mAb (clone HIT3a, BioLegend, cat#300313) at 1 μg/mL and soluble αCD28 mAb (CD28.2, Invitrogen, cat#MA1-20792) at 2 μg/mL, and/or cells were activated with recombinant human IL-12 and IL-18 (R&D Systems), or IL-2 and IL-33, and/or IL1β and IL-23 for 16–20 hours at 37°C in 4% O2, and Brefeldin A was added for the last 4 hours. All cytokines used at 50ng/mL and from Peprotech unless otherwise specified. After stimulation, cells were stained for intracellular cytokine production as described above. For allogeneic stimulation of prenatal CD4 T cells, adult, unrelated CD14^+^ cells were isolated from PBMCs with the CD14 Positive Selection Kit II (STEMCELL Technologies), plated in a 96-well plate at a density of 0.5M CD14^+^ cells/well and allowed to adhere. Non-adherent cells were removed after 30min and adherent cells were incubated with αMHC DP-DQ-DR mAb (BioLegend, clone T<u>ü</u>39) or isotype control (IgG2a, clone MOPC-173, BioLegend) at 10ug/mL for 30 minutes at 37 C. SI CD4^+^ T cells isolated and enriched as above, and stained with Cell Trace Violet (CTV; Invitrogen) to differentiate these from residual adult T cells and co-cultured at a 4:1 (T:APC) cell ratio for 16 hours, with Brefeldin A added in the last 4 hours. Cells were stained for intracellular CD154, and cytokine production as described above.

### Naïve CD4 T cell expansion, maturation, and differentiation assays

Mature CD4 single positive (SP) thymocytes were isolated as above and served as the source of naïve CD4 T cells. Cells were cultured with IL-7 (5 ng/mL), IL-2 (10 ng/mL), or IL-15 (10 ng/mL) for 7 days with media changes every 2-3 days. Cells were kept at 37C and 4% O2 for the duration of the stimulation. In some cases, IL-7 doses were titrated as indicated, and/or TGFβ (10ng/mL) was added for the duration of culture. TCR stimulation was provided by plate-bound αCD3 (clone HIT3a, BioLegend, cat#300313) at 1μg/mL unless otherwise specified, and soluble αCD28 (clone CD28.2, BD, cat#555725) at 2μg/mL. For naïve T cell maturation and differentiation assays, cells were washed and moved to a new 96-well plate at day 5-6, rested for 24hrs in the absence of cytokines, and either matured further with IL-7 (5ng/mL) and/or IL-2 (10ng/mL) in the presence or absence of TCR stimulation as described above or under Th-skewing conditions for an additional 7 days. Cells were differentiated in the presence of TCR stimulation and IL-2 (10ng/mL) for Th0, with the addition of IL-12 (2.5ng/mL) for Th1, and the addition of IL-1b (10ng/mL) and IL-23 (10ng/mL) for Th17, in the presence or absence of IL-7 (5ng/mL). For Th17 differentiation, cells were cultured in IMDM + 10% FBS with 1X non-essential amino acids, 2mM L-glutamine, 10mM HEPES, and Penicillin/Streptomycin. At day 14 of culture, cells were washed, replated, and rested for 24 hours as above, and re-stimulated with plate-bound αCD3 (1μg/mL) and soluble αCD28 (2μg/mL) or rhIL-12and rhIL-18 (50ng/mL) for 24 hours. Where indicated, cells were re-stimulated with Phorbol 12-myristate 13-acetate (PMA) and Ionomycin for 4 hours. Brefeldin A was added for the last 4 hours of each stimulation.

### Chemical Inhibition

Cells were labelled with CTV to track cell proliferation and cultured with IL-7 (10ng/mL) in the presence or absence of chemical inhibitors at 37C in 4%O2 and 5%CO2. Chemical inhibitors from Selleck Chemicals included 5μM SH-4-54 (cat#S7337), 500nM PD0325901 (StemRD, cat#PD-010), 1uM Pictilisib/GDC-0941 (cat#S1065), 5uM Stat3-in-1 (cat#S0818), 50μM JANEX-1 (cat#S5903), and 10μM Solcitinib (cat#S5917). DMSO vehicle control was used at 1μg/mL and 10μg/mL.

### Protein Extraction and Cytokine Level Quantification from Solid Tissue

Sections of tissue (∼10mg) were taken from the SI lamina propria and MLN and placed in chilled Lysis Buffer (1μL/mL Protease inhibitor, 3μL/mL 500mM PMSF in RIPA Buffer). Tissue was disrupted by mixing 20x with a 1mL pipette tip cut to a 2mm opening then vortexed for 15 seconds. Tubes were shaken at 4C for 20 minutes at 300rpm and then the entire process was repeated. Tissue was centrifuged at 14,000 rpm for 10 minutes at 4C and supernatant was collected. Tissue IL-7 and IL-15 concentrations were measured via Cytokine Bead Array using the Human Hematopoietic Stem Cell Panel Kit (LEGENDplex). IL-12p70, IL-18, IL-1b, and IL-23 concentrations were measured using the Human Inflammation Panel 1 Kit (LEGENDplex).

### Statistics

Data were analyzed using Wilcoxon’s test for paired nonparametric data, and one-way Anova with post-hoc Tukey HSD test for comparison of 3 or more groups. Box plot upper and lower hinges represent the first and third quartiles, the center line indicates the median, and the whiskers extends from the hinge to the highest and lowest value no further than 1.5 * (inter-quartile range) from the hinge unless otherwise stated.

## Study Approval

PBMCs we isolated from Trima residues from Trima Apheresis collection kits and were obtained from healthy donors after written informed consent at the Blood Centers of the Pacific. Human prenatal tissues (16 to 22 weeks gestational age) were obtained from terminations of pregnancy after maternal written informed consent with approval from and under the guidelines of the UCSF Research Protection program. Samples were excluded in the cases of known maternal infection, intrauterine prenatal demise, and/or known or suspected chromosomal abnormality.

## Author Contributions

VL and JH designed research studies. VL, SP, SM, and JH conducted experiments. VL, SP, SM, JH acquired data. VL, SP, DB, GF, and JH analyzed data. VL and JH wrote the manuscript. TB and JH provided material.

## Acknowledgements

This work was supported by the UCSF Clinical and Translational Science Institute (CTSI) Pilot Award for Basic and Translational Investigators (2014908), the NIAID (K08 AI128007), and the Burroughs Wellcome Fund (1019828). We acknowledge the Parnassus Flow Cytometry CoLab (PFCC) (RRID:*SCR_018206*) for assistance generating Flow Cytometry data. Research reported here was supported in part by the DRC Center Grant NIH P30 DK063720.”

## Notes

The authors have declared that no conflict of interest exists.

### Competing Interest Statement

The authors have declared no competing interest.

### Summary of Updates

added orchid ID, adjusted figure size

## References

1. Spencer J, Dillon SB, Isaacson PG, Macdonald TT. T cell subclasses in fetal human ileum. Clin Exp Immunol. 1986;65(3):553–8.

2. Lobach DF, Haynes BF. Ontogeny of the human thymus during fetal development. J Clin Immunol. 1987 Mar;7(2):81–97.

3. Darrasse-Jeze G. Ontogeny of CD4+CD25+ regulatory/suppressor T cells in human fetuses. Blood. 2005;105(12):4715–21.

4. Cupedo T, Nagasawa M, Weijer K, Blom B, Spits H. Development and activation of regulatory T cells in the human fetus. Eur J Immunol. 2005 Feb;35(2):383–90.

5. Jain N. The early life education of the immune system: Moms, microbes and (missed) opportunities. Gut Microbes. 2020 Nov 9;12(1):1824564.

6. Rudd BD. Neonatal T Cells: A Reinterpretation. Annu Rev Immunol. 2020 Apr 26;38(1):229–47.

7. Zhu J, Yamane H, Paul WE. Differentiation of Effector CD4 T Cell Populations. Annu Rev Immunol. 2010 Mar 1;28(1):445–89.

8. Kawabe T, Jankovic D, Kawabe S, Huang Y, Lee PH, Yamane H, et al. Memory-phenotype CD4 + T cells spontaneously generated under steady-state conditions exert innate T H 1-like effector function. Sci Immunol [Internet]. 2017 Jun 9 [cited 2022 Jun 30];2(12). Available from: https://www.science.org/doi/10.1126/sciimmunol.aam9304

9. Min B, McHugh R, Sempowski GD, Mackall C, Foucras G, Paul WE. Neonates Support Lymphopenia-Induced Proliferation. Immunity. 2003 Jan;18(1):131–40.

10. Smith NL, Patel RK, Reynaldi A, Grenier JK, Wang J, Watson NB, et al. Developmental Origin Governs CD8+ T Cell Fate Decisions during Infection. Cell. 2018 Jun;174(1):117-130.e14.

11. Masopust D, Vezys V, Marzo AL, Lefrançois L. Preferential Localization of Effector Memory Cells in Nonlymphoid Tissue. Science. 2001 Mar 23;291(5512):2413–7.

12. Sathaliyawala T, Kubota M, Yudanin N, Turner D, Camp P, Thome JJC, et al. Distribution and Compartmentalization of Human Circulating and Tissue-Resident Memory T Cell Subsets. Immunity. 2013 Jan;38(1):187–97.

13. Schreurs RRCE, Baumdick ME, Sagebiel AF, Kaufmann M, Mokry M, Klarenbeek PL, et al. Human Fetal TNF-α-Cytokine-Producing CD4+ Effector Memory T Cells Promote Intestinal Development and Mediate Inflammation Early in Life. Immunity. 2019 Feb;50(2):462-476.e8.

14. Halkias J, Rackaityte E, Hillman SL, Aran D, Mendoza VF, Marshall LR, et al. CD161 contributes to prenatal immune suppression of IFN-γ–producing PLZF+ T cells. J Clin Invest. 2019 Jul 29;129(9):3562–77.

15. Stras SF, Werner L, Konnikova L. Maturation of the Human Intestinal Immune System Occurs Early in Fetal Development. Dev Cell. 2019;51:357–73.

16. Li N, van Unen V, Abdelaal T, Guo N, Kasatskaya SA, Ladell K, et al. Memory CD4+ T cells are generated in the human fetal intestine. Nat Immunol. 2019 Mar;20(3):301–12.

17. Azizi E, Carr A, Plitas G, Rudensky A, Pe’er D. Single-Cell Map of Diverse Immune Phenotypes in the Breast Tumor Microenvironment. Cell. 2018;(174):1293–308.

18. Szabo PA, Levitin HM, Miron M, Snyder ME, Senda T, Yuan J, et al. Single-cell transcriptomics of human T cells reveals tissue and activation signatures in health and disease. Nat Commun. 2019 Dec;10(1):4706.

19. Low JS, Farsakoglu Y, Amezcua Vesely MC, Sefik E, Kelly JB, Harman CCD, et al. Tissue-resident memory T cell reactivation by diverse antigen-presenting cells imparts distinct functional responses. J Exp Med. 2020 Aug 3;217(8):e20192291.

20. Simoni Y, Becht E, Fehlings M, Loh CY, Koo SL, Teng KWW, et al. Bystander CD8+ T cells are abundant and phenotypically distinct in human tumour infiltrates. Nature. 2018 May;557(7706):575–9.

21. Alanio C, Nicoli F, Sultanik P, Flecken T, Perot B, Duffy D, et al. Bystander hyperactivation of preimmune CD8+ T cells in chronic HCV patients. eLife. 2015 Nov 14;4:e07916.

22. Tough DF, Borrow P, Sprent J. Induction of Bystander T Cell Proliferation by Viruses and Type I Interferon In Vivo. Sci New Ser. 1996;272(5270):1947–50.

23. Lee HG, Cho MZ, Choi JM. Bystander CD4+ T cells: crossroads between innate and adaptive immunity. Exp Mol Med. 2020 Aug;52(8):1255–63.

24. Fry TJ, Connick E, Falloon J, Lederman MM, Liewehr DJ, Spritzler J, et al. A potential role for interleukin-7 in T-cell homeostasis. 2001;97(10):8.

25. Kassiotis G, Zamoyska R, Stockinger B. Involvement of Avidity for Major Histocompatibility Complex in Homeostasis of Naive and Memory T Cells. J Exp Med. 2003 Apr 21;197(8):1007–16.

26. Palmer MJ, Mahajan VS, Chen J, Irvine DJ, Lauffenburger DA. Signaling thresholds govern heterogeneity in IL-7-receptor-mediated responses of naïve CD8 + T cells. Immunol Cell Biol. 2011 Jul;89(5):581–94.

27. Sood A, Lebel M, Dong M, Fournier M, Vobecky SJ, Haddad É, et al. CD5 levels define functionally heterogeneous populations of naïve human CD4 + T cells. Eur J Immunol. 2021 Jun;51(6):1365–76.

28. Michaëlsson J, Mold JE, McCune JM, Nixon DF. Regulation of T Cell Responses in the Developing Human Fetus. J Immunol. 2006 May 15;176(10):5741–8.

29. Mold JE, Michaëlsson J, Burt TD, Muench MO, Beckerman KP, Busch MP, et al. Maternal Alloantigens Promote the Development of Tolerogenic Fetal Regulatory T Cells in Utero. Science. 2008 Dec 5;322(5907):1562–5.

30. Schönland SO, Zimmer JK, Lopez-Benitez CM, Widmann T, Ramin KD, Goronzy JJ, et al. Homeostatic control of T-cell generation in neonates. Blood. 2003 Aug 15;102(4):1428–34.

31. Barata JT, Durum SK, Seddon B. Flip the coin: IL-7 and IL-7R in health and disease. Nat Immunol. 2019 Dec;20(12):1584–93.

32. Seddon B, Zamoyska R. TCR and IL-7 Receptor Signals Can Operate Independently or Synergize to Promote Lymphopenia-Induced Expansion of Naive T Cells. J Immunol. 2002 Oct 1;169(7):3752–9.

33. Chattopadhyay PK, Yu J, Roederer M. Live-cell assay to detect antigen-specific CD4+ T-cell responses by CD154 expression. Nat Protoc. 2006 Jun;1(1):1–6.

34. Lee YJ, Jeon YK, Kang BH, Chung DH, Park CG, Shin HY, et al. Generation of PLZF+ CD4+ T cells via MHC class II–dependent thymocyte–thymocyte interaction is a physiological process in humans. J Exp Med. 2010 Jan 18;207(1):237–46.

35. Sallusto F, Lenig D, Lanzavecchia A. Two subsets of memory T lymphocytes with distinct homing potentials and effector functions. Nature. 1999;401:708–12.

36. Surh CD, Sprent J. Homeostasis of Naive and Memory T Cells. Immunity. 2008 Dec;29(6):848–62.

37. Bains I, Antia R, Callard R, Yates AJ. Quantifying the development of the peripheral naive CD4+ T-cell pool in humans. 2009;113(22):5480–7.

38. den Braber I, Mugwagwa T, Vrisekoop N, Westera L, Mögling R, Bregje de Boer A, et al. Maintenance of Peripheral Naive T Cells Is Sustained by Thymus Output in Mice but Not Humans. Immunity. 2012 Feb;36(2):288–97.

39. Tan JT, Dudl E, LeRoy E, Murray R, Sprent J, Weinberg KI, et al. IL-7 is critical for homeostatic proliferation and survival of naïve T cells. Proc Natl Acad Sci. 2001 Jul 17;98(15):8732–7.

40. Younes SA, Punkosdy G, Caucheteux S, Chen T, Grossman Z, Paul WE. Memory Phenotype CD4 T Cells Undergoing Rapid, Nonburst-Like, Cytokine-Driven Proliferation Can Be Distinguished from Antigen-Experienced Memory Cells. Marrack P, editor. PLoS Biol. 2011 Oct 11;9(10):e1001171.

41. Huang HY, Luther SA. Expression and function of interleukin-7 in secondary and tertiary lymphoid organs. Semin Immunol. 2012 Jun;24(3):175–89.

42. Rackaityte E, Halkias J, Fukui EM, Mendoza VF, Hayzelden C, Crawford ED, et al. Viable bacterial colonization is highly limited in the human intestine in utero. Nat Med. 2020 Apr;26(4):599–607.

43. Mishra A, Lai GC, Yao LJ, Aung TT, Shental N, Rotter-Maskowitz A, et al. Microbial exposure during early human development primes fetal immune cells. Cell. 2021 Jun;184(13):3394–409.

44. Seddon B, Legname G, Tomlinson P, Zamoyska R. Long-Term Survival but Impaired Homeostatic Proliferation of Naïve T Cells in the Absence of p56lck. Sci New Ser. 2000;290(5489):127–31.

45. Kawabe T, Yi J, Kawajiri A, Hilligan K, Fang D, Ishii N, et al. Requirements for the differentiation of innate T-bet high memory-phenotype CD4+ T lymphocytes under steady state. Nat Commun. 2020 Dec;11(1):3366.

46. Hirota K, Turner JE, Villa M, Duarte JH, Demengeot J, Steinmetz OM, et al. Plasticity of TH17 cells in Peyer’s patches is responsible for the induction of T cell–dependent IgA responses. Nat Immunol. 2013 Apr;14(4):372–9.

47. Ivanov II, McKenzie BS, Zhou L, Tadokoro CE, Lepelley A, Lafaille JJ, et al. The Orphan Nuclear Receptor RORγt Directs the Differentiation Program of Proinflammatory IL-17+ T Helper Cells. Cell. 2006 Sep;126(6):1121–33.

48. Stockinger B, Omenetti S. The dichotomous nature of T helper 17 cells. Nat Rev Immunol. 2017 Sep;17(9):535–44.

49. McGovern N, Shin A, Low G, Low D, Duan K, Yao LJ, et al. Human fetal dendritic cells promote prenatal T-cell immune suppression through arginase-2. Nature. 2017 Jun;546(7660):662–6.

50. May K, Grube M, Malhotra I, Long CA, Singh S, Mandaliya K, et al. Antibody-Dependent Transplacental Transfer of Malaria Blood-Stage Antigen Using a Human Ex Vivo Placental Perfusion Model. PLoS ONE. 2009 Nov 24;4(11):e7986.

51. Halkias J, Melichar HJ, Taylor KT, Ross JO, Yen B, Cooper SB, et al. Opposing chemokine gradients control human thymocyte migration in situ. J Clin Invest. 2013 May 1;123(5):2131–42.

52. Dobin A, Davis CA, Schlesinger F, Drenkow J, Zaleski C, Jha S, et al. STAR: ultrafast universal RNA-seq aligner. Bioinformatics. 2013 Jan;29(1):15–21.

53. Anders S, Huber W. Differential expression analysis for sequence count data. Genome Biol. 2010;R106.

54. Finak G, McDavid A, Yajima M, Deng J, Gersuk V, Shalek AK, et al. MAST: a flexible statistical framework for assessing transcriptional changes and characterizing heterogeneity in single-cell RNA sequencing data. Genome Biol. 2015 Dec;16(1):278.

55. Bunis DG, Andrews J, Fragiadakis GK, Burt TD, Sirota M. dittoSeq: universal user-friendly single-cell and bulk RNA sequencing visualization toolkit. Alfonso V, editor. Bioinformatics. 2021 Apr 1;36(22–23):5535–6.

56. Zhou Y, Zhou B, Pache L, Chang M, Khodabakhshi AH, Tanaseichuk O, et al. Metascape provides a biologist-oriented resource for the analysis of systems-level datasets. Nat Commun. 2019 Dec;10(1):1523.

