## Supplemental Material for "Homeostatic cytokines reciprocally modulate the emergence of prenatal effector PLZF+CD4+ T cells in humans" (from another version of this preprint: version 1, 2022-09-19)

**Supplementary Information**

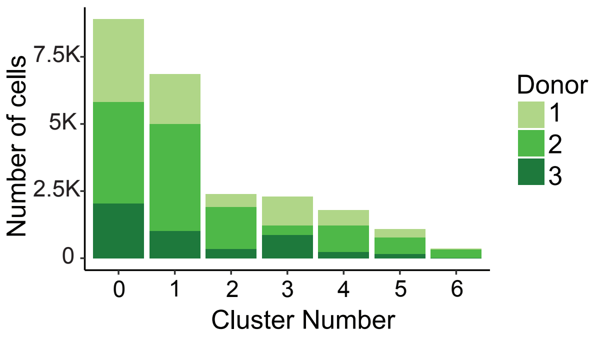


**Figure S1:** Number of cells contributed by donor in each cluster derived from the transcriptional profile of 23,676 individual PLZF$^+$ Vα7.2$^-$ CD4$^+$ TCRαβ$^+$ T cells.

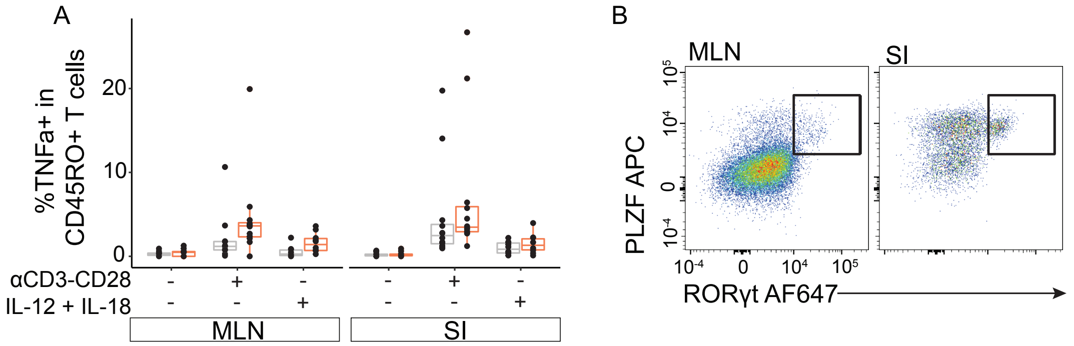


**Figure S2: Prenatal PLZF$^+$ CD4$^+$ T cells are a functionally heterogeneous population.**

(A) Quantification of TNFα$^+$ frequencies after indicated stimulation conditions among PLZF$^-$ (grey) and PLZF$^+$(orange) CD45RO$^+$ CD4$^+$ T cells isolated from the small intestine (SI) and mesenteric lymph node (MLN). (B) Representative flow plots of intracellular RORγt expression of CD4$^+$ T cells isolated from the mesenteric lymph node (MLN) and the small intestine (SI). Circles represent individual donors.

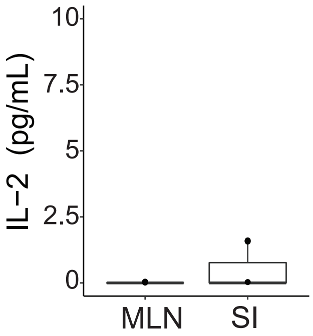


**Figure S3:** Quantification of whole tissue concentration of IL-2 in the prenatal MLN and SI measured by cytokine bead array. Circles represent individual donors. Wilcoxon Signed Rank Test *p < 0.05, **p < 0.01, ***p<0.001 ****p < 0.0001.

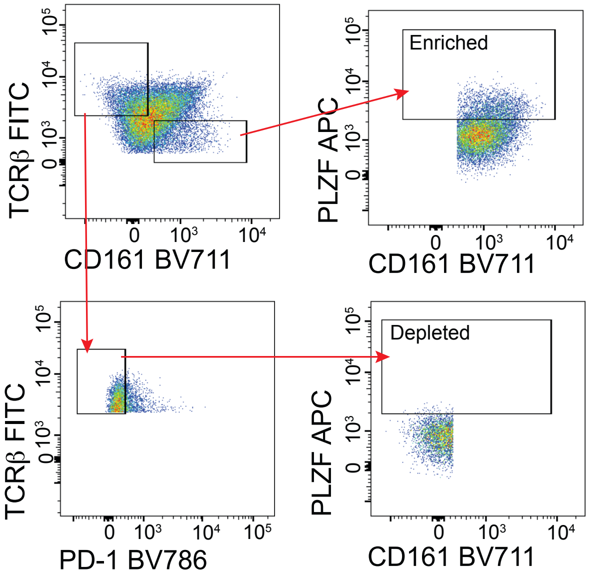


**Figure S4:** Sorting strategy for enrichment and depletion of $PLZF^+$ cells amongst $CD4^+$ naïve T cells based on indicated proxy markers.

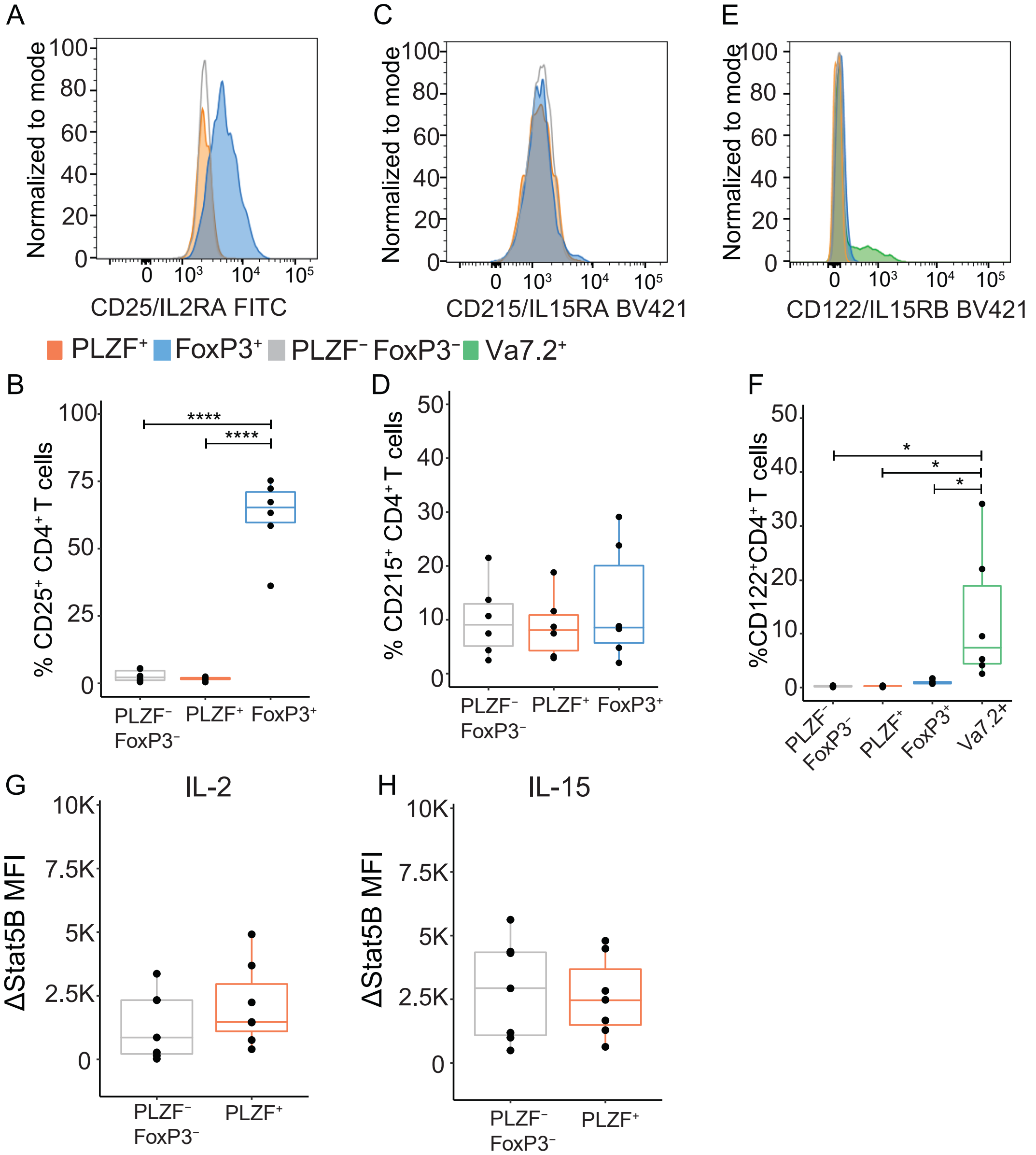


**Figure S5: IL-2 and IL-15 signaling is comparable across prenatal naïve CD4$^+$ T cells.**

(A, C, E) Representative histograms and (B, D, F) frequencies of CD25, CD215, and CD122 expression within indicated populations of naïve CD4$^+$ T cells. (G, H) Normalized MFI of pSTAT5B phosphorylation in prenatal PLZF$^+$ or PLZF$^-$ CD4 naïve T cells after stimulation for 30 minutes with (G) IL-2 or (H) IL-15. Circles represent individual donors. Paired ANOVA with Tukey's multiple comparison test (B, D, F) and Wilcoxon Signed Rank Test (G, H) *p < 0.05, **p < 0.01, ***p<0.001 ****p < 0.0001.

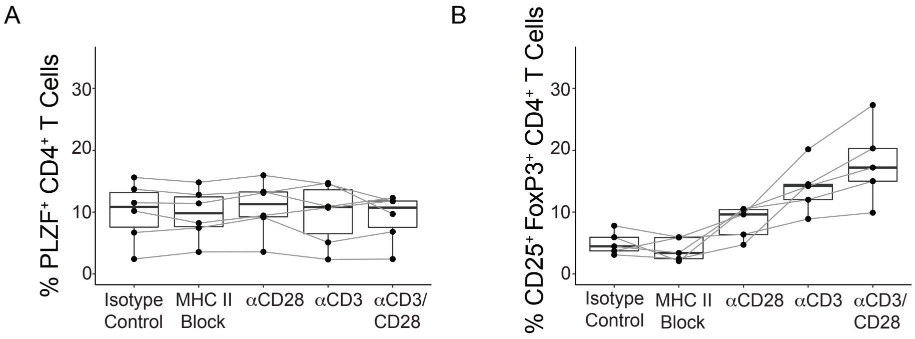


**Figure S6: TCR signaling does not impede IL-7 driven expansion of PLZF$^{+}$CD4$^{+}$ T cells.**

Proportion of (A) PLZF$^{+}$ T cells generated in the presence of IL-7 and (B) CD25$^{+}$FoxP3$^{+}$ cells generated in the presence of IL-2 from culture of naïve CD4$^{+}$ T cells for 7 days in indicated conditions. Circles represent individual donors.

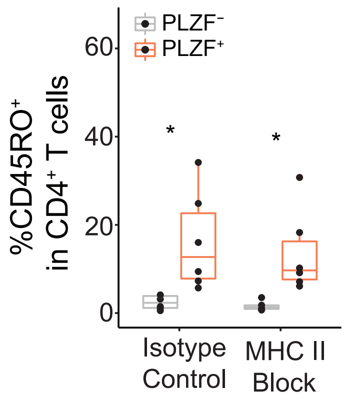


**Figure S7: Emergence of CD45RO$^{+}$ CD4$^{+}$ T cells is unaffected by MHC II Blockade.** Circles represent individual donors.

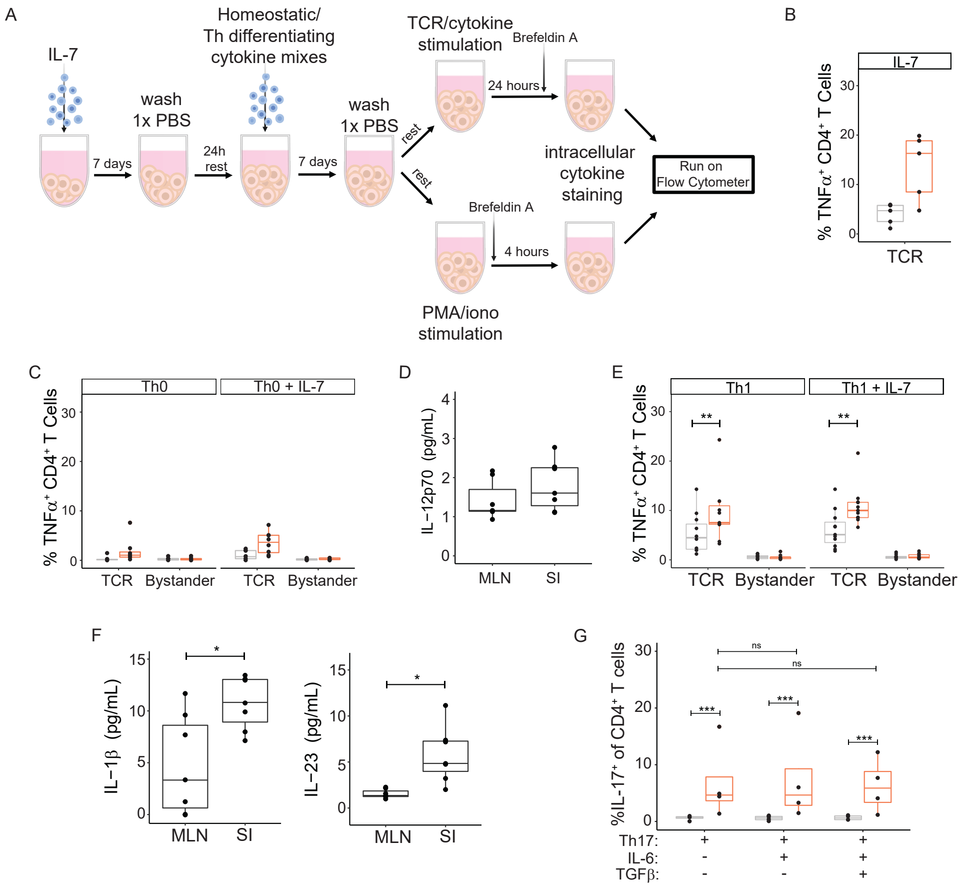


**Figure S8: Tissue-specific cytokines contribute to prenatal $CD4^+$ T cell maturation.**

(A) Schema of the two-step in vitro culture system used in Figure 7A-F. (B,C,E,G) Functional studies following a two-step in vitro culture in which naïve $CD4^+$ T cells were first expanded in the presence of IL-7 alone, followed by 7 days of maturation in the indicated conditions: IL-7 alone, IL-7 + TCR (in the presence of $\alpha$CD3/CD28), Th0 (TCR + IL-2); Th1 (TCR + IL-12), Th17 (TCR + IL-23 + IL-1$\beta$). Proportions of (B, C, E) $TNF\alpha^+$ and (G) IL-$17^+$ cells among $CD4^+$ T cells matured in the indicated conditions following 24hour re-stimulation with $\alpha$CD3/CD28 (TCR) or IL-12/IL-18 (Bystander). Brefeldin A was added in the final 4 hours of all re-stimulations. Quantification of whole tissue concentrations of (D) IL-12p70, and (F) IL-1$\beta$ and IL-23 in the prenatal MLN and SI measured by cytokine bead array. Circles represent individual donors. Wilcoxon Signed Rank Test (B-G) *$p < 0.05$, **$p < 0.01$, ***$p<0.001$ ****$p < 0.0001$.
